# Disrupted Sleep-Dependent Neural Oscillations and Transcriptomic Changes in the *Cacna1g* Loss-of-Function Mouse Model Implicated in Schizophrenia

**DOI:** 10.64898/2026.02.04.703851

**Authors:** Ahmet S. Asan, Nataliia Kozhemiako, Nathaniel Goble, Todd Yau, Tony T. Zhu, Sean K. Simmons, Nathan Haywood, Yining Wang, Soyeon Kim, Zhenglin Guo, Eunah Yu, Soonwook Choi, Lingling Yang, Joshua Z. Levin, Hailiang Huang, Shaun M Purcell, Raozhou Lin, Jen Q. Pan

## Abstract

Sleep is an essential biological process for maintaining brain function, and impaired sleep microarchitecture is a well-established feature of schizophrenia (SCZ). Rare loss-of-function variants in *CACNA1G*, encoding the T-type calcium channel Ca_V_3.1, confer substantial risk for SCZ, implicating sleep-regulatory circuits in disease pathophysiology. Here, we first show that among individuals with SCZ, carriers of rare *CACNA1G* missense variants exhibit more pronounced alterations in sleep neurophysiology than non-carriers, identifying a human phenotype suggestive of altered channel function.

Motivated by this observation, we examined the consequences of *Cacna1g* loss of function in mice. *Cacna1g* deficiency produced profound disruptions in sleep microarchitecture, including reduced sleep spindles, altered slow oscillations, impaired spindle–slow oscillation coupling, and reduced cortical synchrony—closely recapitulating neurophysiological signatures observed in individuals with schizophrenia. Beyond sleep structure, *Cacna1g* loss disrupted the normal coordination of brain rhythms across the 24-hour cycle, most notably abolishing diurnal modulation of theta oscillations.

At the molecular level, *Cacna1g* loss uncoupled sleep–wake state from gene and protein regulation, producing widespread, sleep-phase–biased transcriptional and synaptic alterations across brain regions and cell types, particularly in cortical excitatory neurons. These network and molecular disruptions were accompanied by behavioral hyperactivity during the active phase.

Together, these findings establish *CACNA1G/*Ca_V_3.1 as a key regulator linking sleep-dependent brain dynamics to molecular homeostasis and behavior, and provide a mechanistic framework through which genetic risk for SCZ may drive disease-relevant sleep and circuit dysfunction.

**One Sentence Summary:** Loss of *CACNA1G* disrupts sleep rhythms and gene regulation, creating a mismatch between brain state and behavior linked to schizophrenia.

## INTRODUCTION

Sleep is an essential for brain health and function (*1*). During sleep, the brain undergoes active maintenance and repair processes, including metabolic waste clearance, circuit stabilization, and the generation of coordinated network oscillations - such as slow oscillations and sleep spindles - that support memory consolidation, stress regulation, and information processing (*2–7*). Sleep disturbances are prevalent across brain disorders including schizophrenia (SCZ), in which patients exhibit altered sleep microstructures characterized by abnormalities in sleep spindle and slow oscillatory activities (*8–14*). Notably, a composite of 12 sleep neurophysiology metrics, including sleep spindles and slow oscillation measures, distinguished case versus control status with an AUC greater than 0.93 (*15*) indicating that sleep neurophysiology, at least in part, recapitulated disease-related pathophysiology. Consistent with these findings, genes associated with sleep-related activity are dysregulated in post-mortem brain tissues from individuals of SCZ (*16, 17*). Collectively, these sleep disturbances are increasingly recognized as not only a contributor but a potential driver in symptom expression and cognitive disruptions (*16–19*). However, animal models that comprehensively recapitulate these neurophysiological impairments observed in patients are lacking, and the molecular mechanisms underlying these disruptions remain poorly understood.

*CACNA1G* encodes the Ca_V_3.1 calcium channel, one of three low-voltage-activated T-type calcium channels that open at more hyperpolarized membrane potentials, inactivate rapidly, and deactivate with much slower kinetics compared to high-voltage-activated calcium channels (*20–22*). These channels mediate low-threshold calcium spiking, underlie thalamic firing, and play a pivotal role in neuronal excitability across various brain regions (*23–27*). Rare pathogenic variants in *CACNA1G* have been identified in patients with ataxia, epilepsy, and neurodevelopmental disorders (*28–33*). More recently, large-scale exome sequencing has demonstrated an enrichment of rare protein-truncating variants in *CACNA1G* among individuals with schizophrenia (SCZ), implicating loss-of-function (LoF) of *CACNA1G* in disease risk (*34*). Previous studies have demonstrated that the loss of *Cacna1g* impairs sleep-dependent brain oscillation and disrupts sleep macro-architecture in mice (*23, 35–38*). Together, these findings implicate *Cacna1g*/Ca_V_3.1 in sleep-wake neurophysiology, and suggest that its genetic association with SCZ provides a mechanistic framework to investigate how sleep disturbances contribute to disease pathophysiology.

Despite the strong genetic and neurophysiological evidence linking *CACNA1G* and thalamocortical circuit function to SCZ, the molecular mechanisms by which Ca_V_3.1 dysfunction gives rise to system-level pathology remain unknown. Specifically, the role of Ca_V_3.1 in coordinating diurnal brain oscillations and maintaining molecular homeostasis has not been examined. To address this gap, we first leveraged clinical EEG data to demonstrate that *CACNA1G* missense variants are associated with pronounced alterations in sleep in individuals with SCZ. We then show that *Cacna1g* loss-of-function (LoF) not only recapitulates the patient-associated phenotypes, including sleep spindle deficits, but also induces a global disruption of diurnal brain oscillations in the theta frequency range. Mechanistically, we demonstrate that this network-level pathology is accompanied by dysregulation of sleep-wake-dependent gene expression, leading to the emergence of a cellular stress signature, potentially mediated by *Xbp1* and other transcription factors, and subsequent remodeling of the synaptic proteome. These molecular and synaptic changes are accompanied by behavioral alterations. Collectively, these findings establish *Cacna1g*/Cav3.1 as a critical regulator of sleep-dependent neural dynamics at both network and molecular levels, and suggest that disrupted sleep microstructure and gene regulation during sleep contribute to SCZ pathophysiology.

## RESULTS

### Missense variants in *CACNA1G* are associated with exacerbated sleep neurophysiological disturbances in individuals with SCZ

To test the hypothesis that *CACNA1G* variants exacerbate the core neurophysiological endophenotype of Schizophrenia (SCZ), we leveraged whole-exome sequencing and electroencephalography (EEG) data from the Global Research Initiative on the Neurophysiology of Schizophrenia (GRINS) cohort. Within this cohort (*8, 15*), 14 individuals (out of 301 individuals) carried *CACNA1G* variants, including 12 diagnosed with SCZ, supporting an association between *CACNA1G* variants and SCZ risk (Fisher’s exact test, OR = 4.6, p = 0.049). We next asked whether these 12 variant-carrying SCZ patients exhibited more pronounced sleep neurophysiology abnormalities – that is, a more strongly “SCZ-like” profile- relative to no carrier SCZ patients. To this end, we first defined an SCZ-vs-CTR template using all nominally significant metrics (N = 2997) spanning fast/slow spindles, slow oscillations (SO), their coupling, and spectral power (0.5–20 Hz) across 57 EEG channels in the full GRINS sample. We then compared the 12 *CACNA1G* variant carriers with the remaining SCZ cases by quantifying the number of metrics whose directionality matched the SCZ-CTR template. Statistical significance was assessed using a permutation-based null distribution generated by randomly sampling 12 SCZ patients from the cohort (vs. the remaining SCZ cases) 3000 times. CACNA1G variants carriers aligned with the SCZ-CTR template significantly more often than expected by chance (mean matches = 2207, null mean = 1299, *p* = 0.005), indicating exaggerated sleep neurophysiology disruptions in these patients (**Fig. 1A**). Figures 1B–D further illustrate that the differences observed between SCZ patients with CACNA1G alterations and those without closely resemble the SCZ–control contrast for selected sleep EEG features.

**Figure 1.**
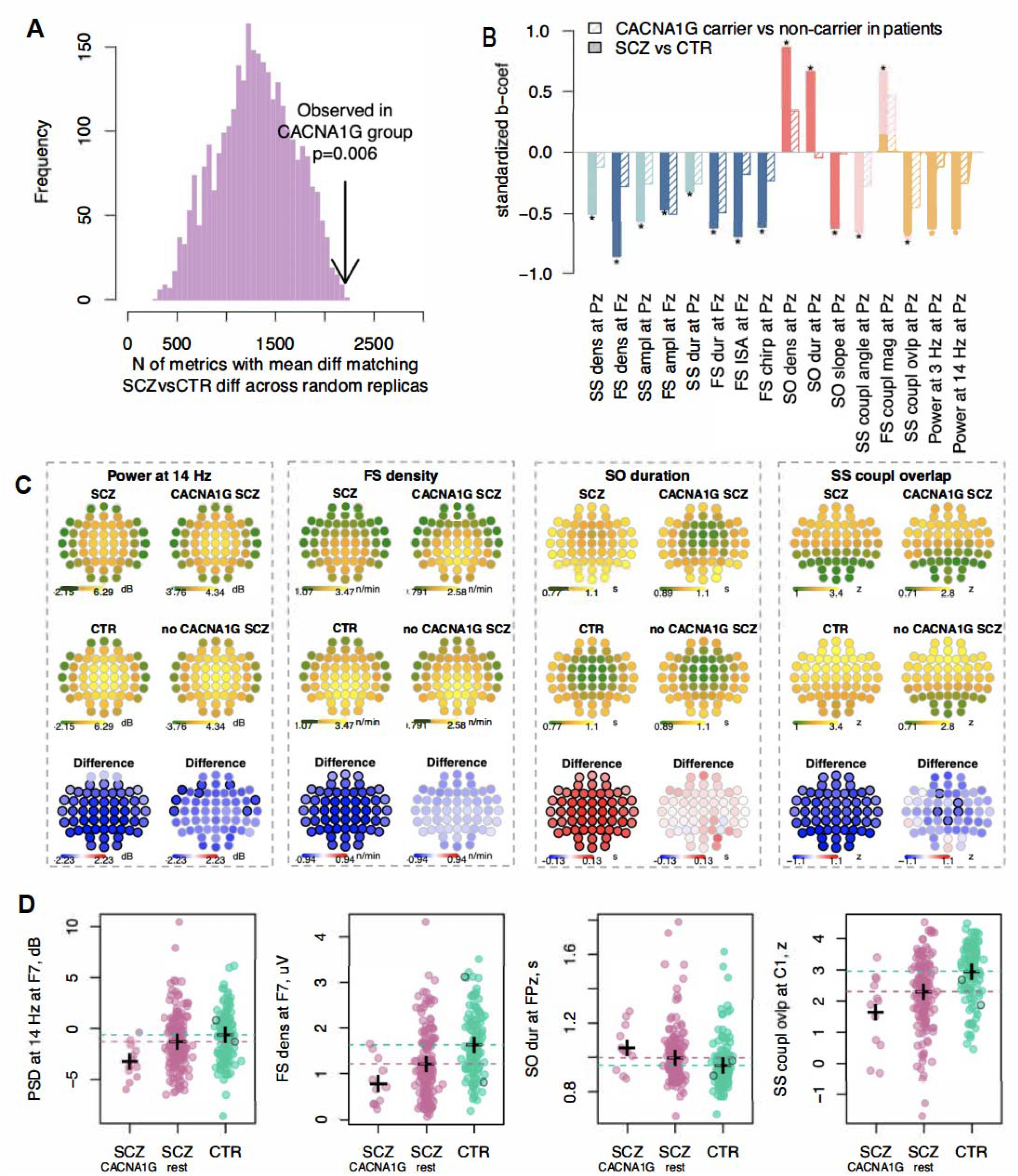
Patients with *CACNA1G* rare missense changes exhibit more severe sleep neurophysiology alterations than other SCZ patients. **(A)** Histogram showing the distribution of metrics matching the SCZ-vs-CTR template across 3,000 random subsamples (12 randomly selected SCZ patients vs. the remaining patients) and the red line represents the number of matches observed in 12 SCZ patients carrying *CACNA1G* missense variants. **(B)** Bar plot showing effect sizes for differences between patient and control individuals (SCZ vs CTR) and differences between carriers and non-carriers of rare *CACNA1G* missense variants among patients (Carrier vs non-carrier), for selected sleep EEG metrics at Fz (frontal) or Pz (parietal) channels highlighting consistent direction compared to SCZ-vs-CTR group differences. Stars indicate nominal significance (p < 0.05) for the respective comparisons. **(C)** Topoplots displaying average metric values and group differences across all channels: SCZ vs. CTR (left), and carrier vs. non-carriers of rare *CACNA1G* missense changes among patients (right) for each panel. Channels with nominally significant differences (p < 0.05) are outlined in bold. **(D)** Scatter plots of exemplar channels showing individual metric values for SCZ patients with *CACNA1G* rare variants (SCZ carriers), SCZ patients without CACNA1G rare variants (SCZ non-carriers), and controls. The two control individuals with *CACNA1G* variants are circled in black. Crosses mark group medians. Horizontal dashed lines indicate medians for SCZ (pink) and CTR (green) groups without rare *CACNA1G* missense variants.

### *Cacna1g* LoF results in altered EEG power spectrum, impaired 24-hour rhythmic patterns, and disrupted cortical connectivity

Motivated by our finding that *CACNA1G* variants exacerbate sleep neurophysiology alterations in individuals with SCZ (**Fig. 1**), we next performed a comprehensive characterization of brain network dynamics in *Cacna1g* loss-of-function (LoF) mice, including heterozygous (*Cacna1g +/-*, HT), homozygous knockout (*Cacna1g -/-*, KO), and wild-type littermates (*Cacna1g +/+*, WT). We recorded continuous 24-hour spontaneous *in vivo* EEG from frontal and parietal cortical regions, as previously described (*39, 40*). We first examined the impact of *Cacna1g* LoF on spontaneous spectral power across frequencies ranging from 0.5 to 50 Hz during the light cycle. Across all genotypes, low-frequency activity (slow oscillations and delta) was higher during NREM sleep compared to wakefulness, while high-frequency activity (e.g., beta, gamma) was more prominent during wakefulness (**Fig. 2B-C**). Despite these shared state-dependent patterns, we identified pronounced genotype-specific differences during both NREM and wake, predominantly in the frontal cortex (**Fig. 2C**). Notably, power of higher frequency oscillations, including beta and gamma bands, was persistently elevated during both wake and NREM in *Cacna1g* HT and KO animals, compared with WT littermates. Conversely, power in lower frequency oscillations, including theta and delta bands, was reduced during wake and NREM, with the exception of frontal delta power during NREM. To further assess spindle-related activity, we examined oscillatory power in the 7–20 Hz range encompassing the sigma band. This activity decreased in HT and KO animals and showed significant group differences during NREM in both regions (**Fig. 2C, inset**). These power changes during the light cycle were not significantly altered in the parietal cortex (**Fig. 2C**). Interestingly, during the dark cycle, the parietal region also showed significant changes during NREM, highlighting region- and period-specific variations (Fig. S1).

**Figure 2.**
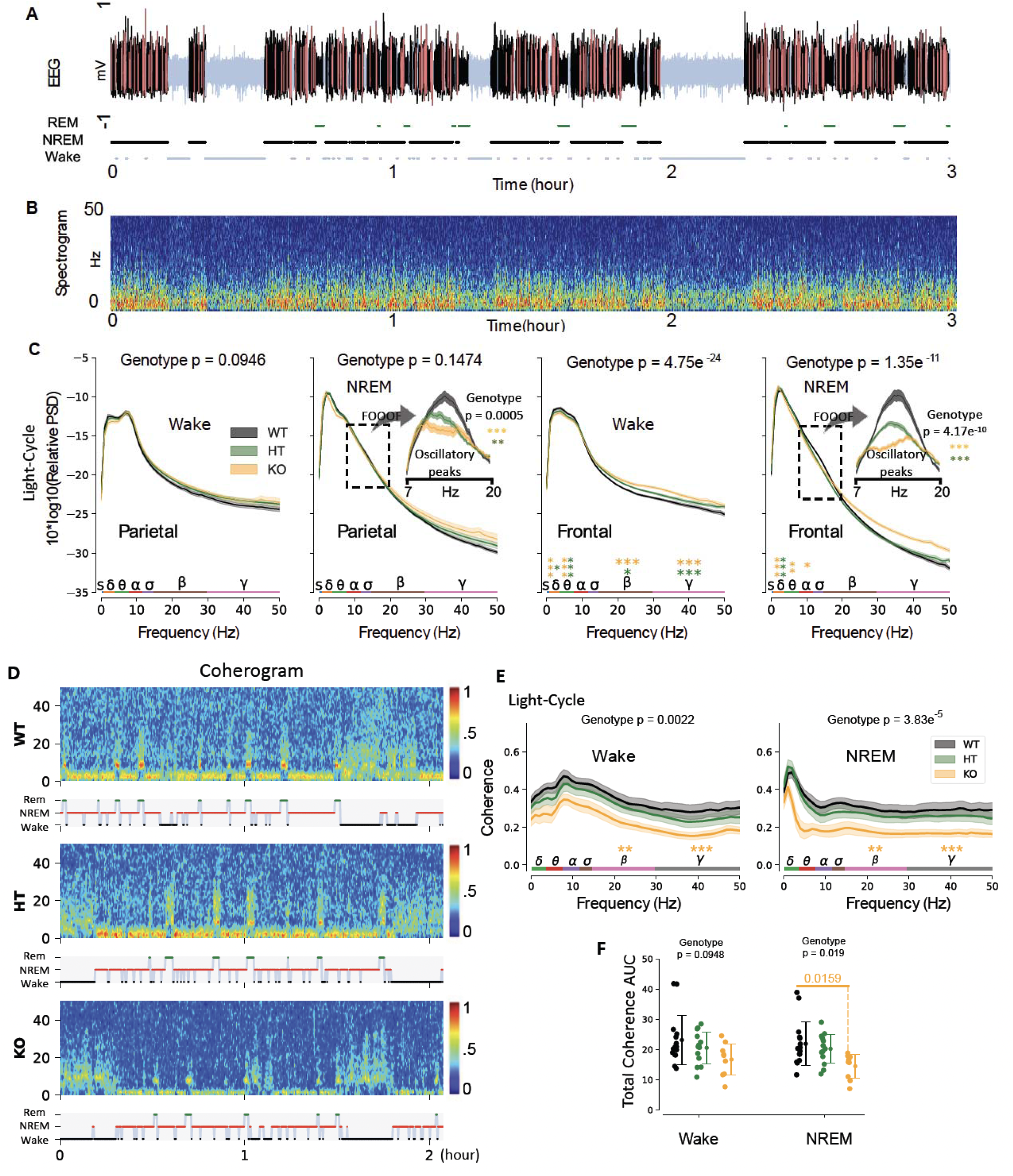
*Cacna1g* Loss of Function Alters Cortical Oscillations and Disrupts Cortical Connectivity. **(A)** Representative raw EEG recording: light blue indicates wakefulness, black represents sleep, and red marks sleep spindles. The hypnogram below shows the corresponding vigilance states: wake (black), REM (green), and NREM (light blue). **(B)** Spectrogram of the EEG recording, displaying frequency content over time. **(C)** Power spectral density comparison during wakefulness and NREM sleep. Thick lines indicate group means, and shaded areas represent standard error. Orange and green asterisks mark significant differences for KO and HT compared to WT littermates in specific frequency bands, respectively. The insets highlight the periodic component, extracted using the FOOOF algorithm, within the dashed window of 7-20 Hz. Statistical significance: *p < 0.05, **p < 0.01, *p < 0.001 (linear mixed-effects model followed by Tukey’s multiple comparisons test). **(D)** Representative coherograms illustrating temporal variations in coherence across different genotype groups, revealing reduced overall coherence in a KO mouse compared to WT and HT. **(E)** Averaged coherence spectra across genotype groups, with thick lines representing the mean and shaded areas denoting the standard error. Orange asterisks indicate significant pairwise differences between WT and KO in the beta and gamma bands. Statistical significance: *p < 0.05, **p < 0.01, *p < 0.001 (linear mixed-effects model followed by Tukey’s multiple comparisons test). **(F)** KO animals display reduced total coherence during wakefulness and NREM sleep (*p < 0.05, mean ± standard deviation is shown).

Brain oscillations promote synchronization across cortical regions, enabling coordinated information processing. To understand how disruptions in these oscillations affect cortical communication, we examined connectivity between the two recorded brain regions. As shown in sample recordings from each genotype, NREM sleep was associated with increased delta coherence, whereas faster frequencies such as alpha and beta were elevated during wakefulness and REM sleep across all groups (**Fig. 2D**). Group-level comparison revealed that KO animals showed a robust reduction in the coherence between the frontal and parietal cortex in the beta and gamma bands during both wakefulness (KO vs. WT, beta, *p*□=□0.002; gamma, *p*□<□0.0001) and NREM (KO vs. WT, beta *p*□=□0.0013; gamma, *p*□=□0.0001) during the light cycle, with similar reductions observed in the dark cycle (**Fig. 2E**, S2). Consistent with these frequency-specific deficits, total coherence across the full spectral power was also reduced in KO animals during NREM in both dark and light cycles (KO vs. WT, light *p* = 0.0159; dark, *p* = 0.008, **Fig. 2F)**, indicating a global disruption of cortical synchrony.

### *Cacna1g* LoF altered cardinal oscillatory patterns during NREM sleep

Sleep spindles and slow oscillations (SOs) are fundamental features during NREM sleep and are consistently disrupted in SCZ and other psychiatric disorders (*41–44*). Moreover, the precise coupling between SOs and spindles is important for sleep-dependent memory consolidation (*45, 46*). We therefore performed a detailed analysis of event-level properties of these oscillations and their coupling during NREM in *Cacna1g* LoF animals.

We first detected spindles with a central frequency of 11Hz (*40, 47*) and found that spindle density, amplitude, duration, and integrated spindle activity (ISA), were all significantly disrupted by *Cacna1g* LoF during NREM sleep in a gene-dose-dependent manner (**Fig. 3A-B**) in the frontal cortex. In the parietal cortex, spindle density and duration, but not amplitude, were also reduced albeit to a lesser extent than in the frontal channel (**Fig. 3B**). As spindle activity is observed across a broader frequency range (*48*), we analyzed spindles with central frequencies of 9, 13, and 15 Hz. These analyses, together with analyses performed during the dark cycle, revealed comparable NREM-specific spindle deficits (Fig. S3). Collectively, these findings showed that *Cacna1g* LoF leads to marked reduction in sleep spindle activity, consistent with the decreased oscillatory power observed in the sigma frequency range (**Fig. 2C, inset**).

**Figure 3.**
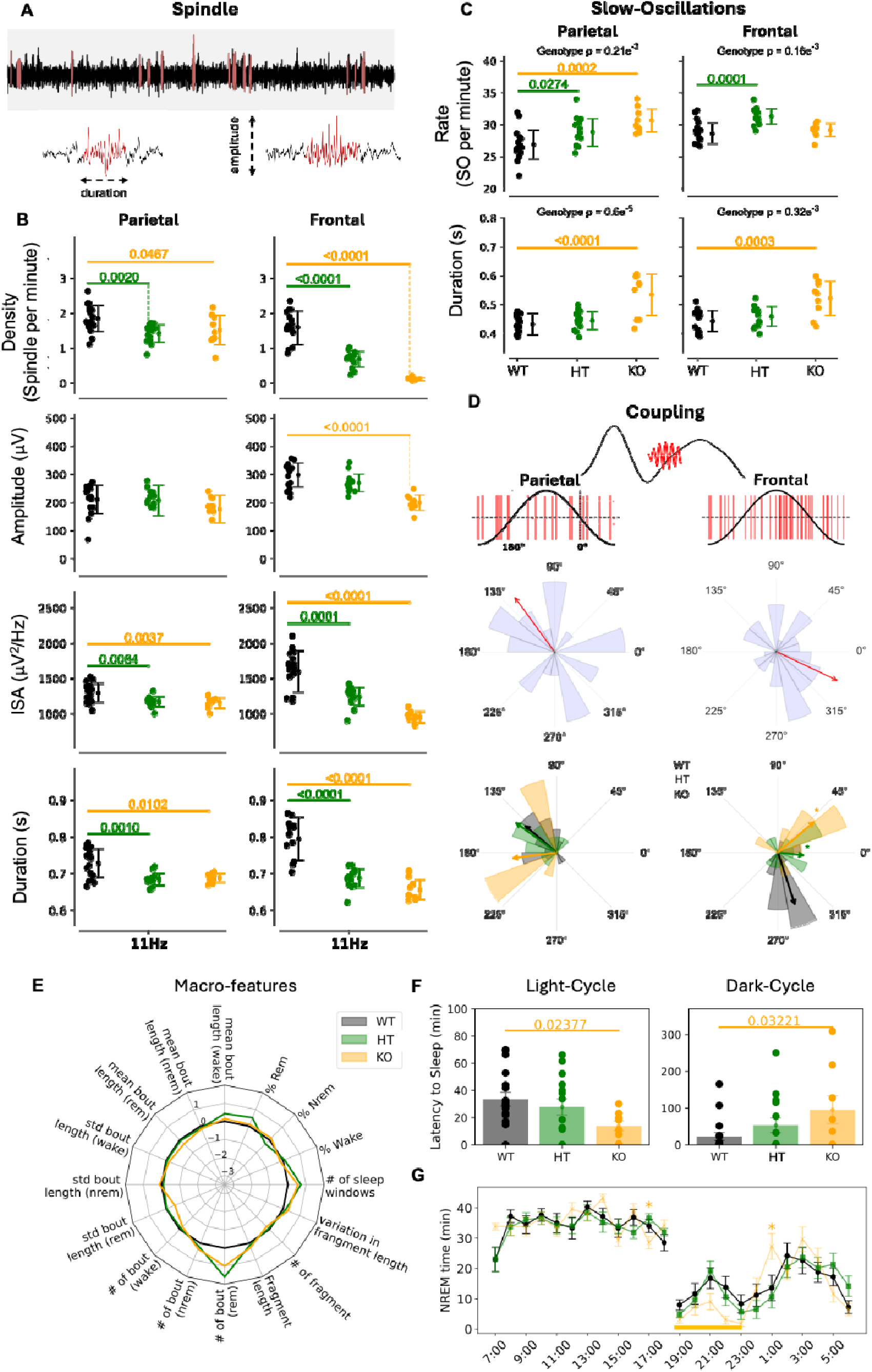
*Cacna1g* Loss of Function Alters Oscillatory Patterns During Sleep in the Light Period. **(A)** A representative recording illustrating detected sleep spindles and their properties. **(B)** Group comparison of spindle density, amplitude, duration, and integrated spindle activity at 11 Hz. Mean ± standard deviation is shown. **(C)** Comparison of slow oscillation rate and duration across genotypes. Mean ± standard deviation. **(D)** Top: Filtered EEG segments illustrating the temporal relationship between slow oscillations and sleep spindles. Middle: Spindle midpoint positions (red lines) overlaid on a schematic slow oscillation (sinusoidal signal), and circular histogram and mean phase direction for a WT animal in parietal and frontal cortical regions. Bottom row: Group-level circular histograms of coupling phase angles with coupling direction and strength indicated by arrows for three genotypes. **(E)** Z-scored comparison of macro sleep features across genotypes. **(F)** Comparison of three genotype groups in latency to sleep during the light and dark cycle. **(G)** NREM duration in 24 hours across three genotype groups. The thick orange line highlights a decreased trend in KO mice relative to WT littermates during the early dark cycle hours.

We next examined SOs during NREM sleep in the light cycle and found that both HT (parietal *p*□=□0.0274; frontal *p*□=□0.0001) and KO (parietal *p*□=□0.0002) animals exhibited a significant increase in SO density relative to their WT littermates (**Fig. 3C**). In addition, KO animals displayed significantly longer SO duration in the parietal (*p*□<□0.0001) and frontal (*p*□=□0.0003) cortices (**Fig. 3C**), whereas SO duration was unchanged in HT animals. Comparable alternations in SO density and duration were observed during NREM sleep in the dark cycle (Fig. S3).

Finally, we examined the coupling between sleep spindles and SOs during the NREM, focusing on 11 Hz spindles. During the light cycle, we observed a significant reduction in coupling density, defined as the number of spindle-SO couplings events normalized to NREM duration - in the frontal cortex (group comparison, *p* < 0.0001; WT vs. KO, *p* < 0.0001) (Fig. S4C). In contrast, coupling strength, quantified as the magnitude of the mean coupling vector, was increased (group comparison, *p* < 0.0001; WT vs. KO, *p* < 0.0001) in KO animals, indicating more tightly clustered coupling angles (Fig. S4C). Consistent with this shift, the ratio of coupled to uncoupled spindles was also elevated in both frontal and parietal cortices (group comparison, frontal *p* < 0.0001; parietal *p* = 0.0005), suggesting a preferential loss of uncoupled spindle events (Fig. S4C).

Analyses of coupling phase revealed region-specific pattern: in the frontal cortex, spindles preferentially coupled to the downswing of SOs (mean angle = 280 degrees), whereas coupling in the parietal cortex was more broadly distributed and centered near the SO upswing (mean angle = 135 degrees, **Fig. 3D**). To assess cross animal consistency, we compared mean coupling angles and observed a significant genotypic effect in the frontal cortex (group comparison, *p* = 0.0002; WT vs. KO, *p* < 0.0001; WT vs. HT, *p* = 0.042, Bonferroni corrected). We next evaluated coupling strength across animals by testing the non-uniformity of coupling angle distributions. WT animals exhibited strong phase clustering in both frontal (*p* < 0.0001) and parietal (*p* = 0.0046) cortices **(Fig. 3D)**. This clustering was selectively disrupted in HT animals in the frontal cortex (*p* = 0.119, **Fig. 3D**), while remaining intact in the parietal cortex. A similar pattern of coupling dynamics was observed during the dark cycle, with significant genotypic differences again predominantly confined to the frontal cortex (Fig. S4B).

### *Cacna1g* LoF exhibited limited impact on sleep macro-architecture

We next examined the sleep macro-structures, including sleep stage length, bout length, and sleep fragmentation, across HT, KO, and their WT littermates using 24-h EEG recordings. Overall, HT and KO animals spent comparable portion of time in NREM sleep, wakefulness, and REM sleep relative to WT, and exhibited no significant differences in mean bout length, number of bouts for per each vigilance state, or other indices of sleep fragmentation, including sleep window analysis (see Methods, **Fig. 3E**). In contrast, KO animals showed significant disruption in sleep latency, characterized by delayed sleep onset during the dark cycle (*p* = 0.032) and shortened latency during the light cycle (*p* = 0.023). HT animals showed a similar but non-significant trend towards delayed sleep onset during dark (*p* = 0.185) (**Fig. 3F**). Analyses of hourly NREM sleep dynamics revealed a reduced NREM duration in KO animals during the early hours of the dark period; however, no significant differences were detected across genotypes when considered over the full 24-h cycle (**Fig. 3G**).

To further evaluate sleep fragmentation, we quantified brief bouts (< 60 s), which are indicative of unstable sleep. KO animals exhibited a higher number of short NREM bouts, although this increase did not reach statistical significance, while the duration of these bouts was significantly reduced compared with WT animals during the light cycle (Fig. S4E). We next analyzed vigilance state transitions, focusing on transitions involving NREM sleep. REM-to-NREM transitions differed significantly across genotypes, with LoF animals exhibiting more frequent transitions than WT controls (*p* = 0.0042), and NREM-to-REM transitions showed a similar, though non-significant, trend (*p* = 0.0534, Fig. S4F).

In summary, *Cacna1g* LoF animals exhibited shorter NREM bouts and increased NREM-REM transitions, indicative of fragmented sleep. Most notably, we identify gene dose-dependent alterations in sleep spindle and slow oscillation activity, as well as their temporal dynamics and coupling, associated with *Cacna1g* LoF. These changes reflect fundamental impairment in NREM sleep.

### Loss of function of *Cacna1g* disrupts diurnal regulation of brain oscillatory power and gene expression programs

The power of brain oscillations across distinct frequency bands exhibits circadian modulation over the course of the day (*49, 50*). To investigate the contribution of *Cacna1g*/Ca_V_3.1 to these circadian patterns, we modeled the hourly oscillatory power within each frequency band using cosinor function (*51*). In the delta band, mice of all three genotypes displayed robust 24-hour rhythmicity across light and dark periods, with the cosinor model accounting for a substantial portion of the variance (R²: WT = 0.65, HT = 0.71, KO = 0.52). Neither rhythm amplitude nor phase differed significantly between *Cacna1g* KO or HT mice and their WT littermates (**Fig. 4A**). In contrast, theta-band activity showed pronounced genotype-dependent effects. Whereas WT and HT mice showed strong circadian rhythmicity (R²: WT = 0.5, HT = 0.6), KO mice displayed a near-complete loss of 24-hour rhythmic modulation (R^2^: KO = 0.15). This disruption was accompanied by a significant genotype-dependent reduction in rhythm amplitude (*p* = 0.0025) and a moderate phase shift (*p* = 0.0873, **Fig. 4B**). Moreover, phase clustering analyses revealed consistent rhythm timing across animals in WT (*p* < 0.0001) and HT (*p* < 0.0001) groups, whereas KO animals exhibited no significant clustering (*p* = 0.0732), indicating highly variable and disorganized theta rhythms across animals (**Fig. 4B**, right panel). Analysis of additional frequency bands revealed significant alternation in alpha-band exclusively in HT animals. Together, these findings indicate that *Cacna1g*/Ca_V_3.1 not only regulates oscillatory power within specific frequency bands but is also essential for maintaining coherent diurnal modulation of brain rhythms across sleep-wake cycle, with the most pronounced effects observed in the theta band.

**Figure 4.**
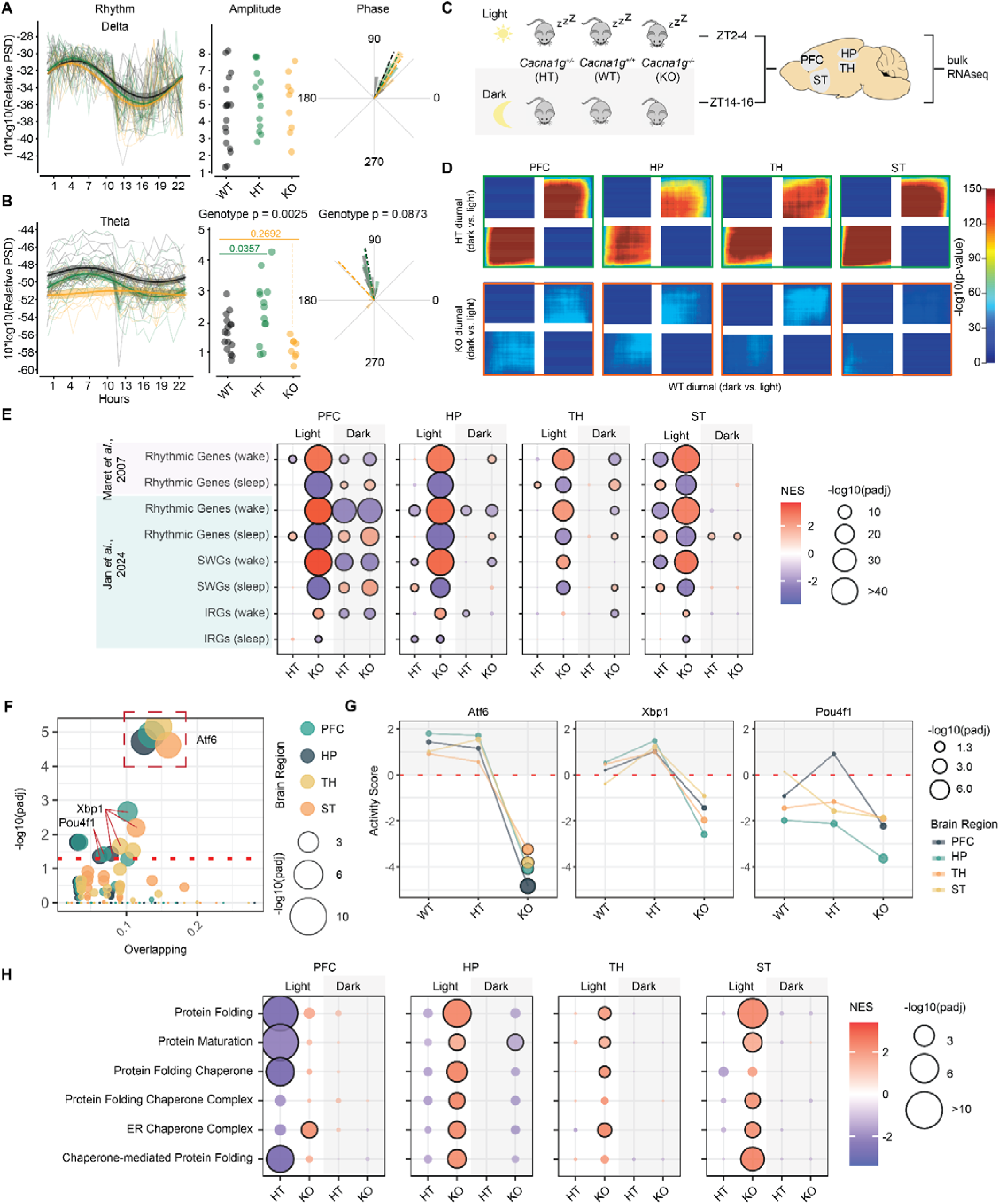
Loss of Function of *Cacna1g* Impairs Diurnal Brain Oscillation and Transcriptomic Changes. **(A, B)** Dynamic changes in delta and theta band power in 24 hours. Thin lines represent individual data, while thick lines indicate fitted cosine curves, with shaded areas representing standard error. Amplitude and phase values derived from the cosine fits are shown in the accompanying plots on the right. **(C)** Experimental design schematic. Wild-type (WT; *Cacna1g*^+/+^), heterozygous (HT; *Cacna1g*^+/–^), and knockout (KO; *Cacna1g*^−/−^) mice were sampled during the light and dark phases across four brain regions: prefrontal cortex (PFC), hippocampus (HP), thalamus (TH), and striatum (ST). At least *N* = 7 mice per genotype per period, except for KO/light (*N* = 3) and KO/dark (*N* = 4). **(D)** Rank-rank hypergeometric overlap (RRHO2) analysis comparing diurnal transcriptomic changes (dark vs. light) in HT and KO mice relative to WT across brain regions. **(E)** Gene set enrichment analysis (GSEA) of circadian rhythmic genes, sleep–wake genes (SWGs), and intrinsic rhythmic genes (IRGs) in HT and KO mice compared to WT. Dot color represents normalized enrichment score (NES), and dot size reflects statistical significance (–log10 adjusted *p* value). Dots outlined in black indicate adjusted *p* < 0.05. **(F)** Transcription factor overrepresentation analysis of wake-associated genes across brain regions. The x-axis shows the overlap ratio (fraction of leading-edge genes within each transcription factor target set), and dot size and y-axis represent enrichment significance (–log10 adjusted *p* value). The dashed red line indicates adjusted *p* = 0.05. **(G)** Inferred transcription factor activity for *Atf6*, *Xbp1*, and *Pou4f1* across genotypes and brain regions based on diurnal gene expression changes. Filled circles denote adjusted *p* < 0.05. **(H)** GSEA of protein folding, maturation, and endoplasmic reticulum (ER)–related pathways in HT and KO mice compared to WT within each light–dark period.

Changes in gene expression and protein regulation across sleep-wake cycles, together with EEG dynamics, provide critical insights into the molecular and neural processes underlying sleep (*52–54*). To complement the observed disruption in brain oscillation dynamics and to uncover the molecular mechanisms associated with sleep disturbances in *Cacna1g* LoF, we performed bulk RNA sequencing (RNA-seq) to profile transcriptional signatures across multiple brain regions in HT, KO, and WT animals. We focused on the prefrontal cortex (PFC), hippocampus (HP), thalamus (TH), and striatum (ST), as these regions are implicated in SCZ pathophysiology, and/or exhibited high expressions of *Cacna1g*/Ca_V_3.1 (Fig. S5A-B). We collected brain tissue during the light cycle (ZT2-4) when the animals were predominantly asleep, and during the dark cycle (ZT14-16) when they were primarily active, enabling evaluation of intrinsic diurnal regulation of the transcriptome (**Fig. 4C**). Principal component analysis (PCA) revealed that samples were clustered primarily by region, rather than by genotype or time point, consistent with known regional transcriptional identity (Fig. S5C). RNA-seq libraries were generated in two independent batches, and a batch effect was evident upon data integration (Fig. S5D). We therefore applied batch-correction procedures (see Methods), which effectively mitigated the batch-associated variance while preserving the major biological signal driven by the diurnal period (Fig. S6B).

To assess if the diurnal signature mirrors the disruption in brain power, we asked if the transcriptomic signature change across periods within WT was different from those changes within HT or KO, respectively. We used rank-rank hypergeometric overlap (RRHO) analysis to highlight shared diurnal transcriptional changes across genotypes. This threshold-free approach enables sensitive detection of concordant transcriptional regulation between conditions. Across all four brain regions, we observed strong concordant overlap between WT and HT mice among both upregulated and downregulated genes, indicating largely preserved diurnal transcriptional programs in HT animals (**Fig. 4D**). In contrast, these concordant signals were markedly attenuated in KO mice (**Fig. 4D**). The pronounced loss of overlap in KO animals directly demonstrated that *Cacna1g* Lof profoundly disrupts normal diurnal regulation of gene expression across all examined brain regions.

### Loss of function of *Cacna1g* causes more profound transcriptomic changes in the light period

To characterize genotype-specific transcriptomic changes across the diurnal cycle, we assessed differential gene expression at each time point and found that *Cacna1g* LoF animals exhibited substantially more differentially expressed genes (DEGs) during the light cycle than during the dark cycle in three of the four brain regions examined (Fig. S6A, table S1, DEGs having adjusted *p* < 0.05). Notably, the thalamus showed the largest number of DEGs in the light cycle. To complement DEG analyses, we applied the TRADE algorithm, which quantifies transcriptomic-wide impact (TI) across genetic perturbations independently of statistical power and experimental context (*55*). Consistent with DEG results, TRADE revealed pronounced transcriptomic disruption during the light cycle, but not dark cycle, across all brain regions in KO animals (Fig. S6A, table S1). PCA further demonstrated clear separation between light- and dark-cycle transcriptomic profiles along both PC1 and PC2 in the prefrontal cortex, thalamus, and striatum. In contrast, such separation was not observed in the hippocampus, suggesting that this region is among the least sensitive regions to period-dependent transcriptional changes in this dataset (Fig. S6B). Consistent with the elevated transcriptomic burden, thalamus also exhibited genotype-dependent separation during the light cycle, highlighting its particular vulnerability to *Cacna1g* LoF (Fig. S6B).

To gain insights into how *Cacna1g* LoF alters brain function, we first performed gene set enrichment analysis (GSEA) using neuronal activity-regulated gene (ARG) sets (*56*), as the expression of these genes, among other implications, can serve as a proxy for measuring recent neuronal activity and encoding long-term memory (*56, 57*). We observed significant enrichment of ARGs among downregulated genes in HT animals during the light period across three of the four brain regions examined (Fig. S6C). In contrast, KO animals exhibited enrichment of ARG expression in the upregulated genes in the thalamus (Fig. S6C, first row). To refine these findings, we next analyzed three mechanistically distinct subsets of ARGs (*56*). Primary response genes (PRGs), which include canonical rapid response genes like *Fos* and *Arc*, were significantly enriched among downregulated genes in the prefrontal cortex, hippocampus, and striatum during the light period in HT animals (Fig. S6C, second row). Delayed PRGs also showed a pronounced reduction in HT animals in striatum in the light, but were unexpectedly increased in KO animals in the thalamus and striatum in the light, potentially due to compensatory mechanisms (Fig. S6C, third row). In contrast, secondary response genes (SRGs) were not significantly changed (Fig. S6C) in any region or condition. Together, these results revealed pronounced effects of *Cacna1g* LoF on activity-dependent transcription during the light period, with HT animals showing broad suppression of ARG programs whereas KO animals exhibited increased ARG expression in the thalamus.

### Loss of function of *Cacna1g* causes changes in multiple molecular pathways across different brain regions in both light and dark periods

To obtain a broader view of the biological processes impacted by *Cacna1g* LoF, we performed GSEA using gene ontology (GO) terms from the molecular signature database (MSigDB) (*58, 59*). Comparisons within the light period between HT and WT animals revealed region specific enrichment patterns: gene sets related to mitochondria functions, such as oxidative phosphorylation and cellular respiration, and to ribosomes, such as cytoplasmic translation, were enriched among downregulated genes in the prefrontal cortex and hippocampus, but among upregulated genes in the thalamus and striatum (Fig. S6D-E, table S2). *Cacna1g* LoF also altered the enrichment of synapse-related gene sets, including those involved in synaptic organization and function (Fig. S6D, table S2). Notably, during the light cycle in HT animals, these synaptic gene sets were enriched among downregulated genes in the prefrontal cortex and hippocampus but among upregulated genes in the striatum, suggesting a regionally divergent effect of *Cacna1g*/Ca_V_3.1 on synaptic programs. Together these findings implicate *Cacna1g*/Ca_V_3.1 in the sleep-dependent modulation of metabolic and synaptic pathways across brain regions, with potential consequences for large scale network connectivity.

Glucocorticoid signaling follows a circadian rhythm, peaking at wake onset, declining throughout the day, and reaching a nadir before sleep, where it plays a key role in regulating period-specific gene expression (*60*). To determine whether glucocorticoid signaling is altered in *Cacna1g* LoF animals, we analyzed gene sets indicative of glucocorticoid receptor (GR) activity. These include transcriptional targets regulated by dexamethasone, a synthetic GR ligand, derived from *in vitro* primary cortical neuron and astrocyte cultures, as well as from hippocampal tissue following *in vivo* acute dosing in mice (table S3). We observed significant enrichment of GR-regulated genes among upregulated genes in both HT and KO animals during the light cycle (Fig. S6F). Given the established role of glucocorticoids in promoting wakefulness and their elevation during sleep deprivation (*61–63*), we next examined gene sets associated with sleep deprivation. Genes regulated by sleep deprivation showed significant overlap with the *Cacna1g* KO signature during the light period. Specifically, both upregulated and downregulated sleep-deprivation markers were directionally consistent with the changes in KO mice (Fig. S6F). Together, these findings indicate that *Cacna1g* KO animals exhibit an altered GR-mediated transcriptional response that partially recapitulates gene expression patterns observed during sleep deprivation, implicating disrupted glucocorticoid signaling as a contributor to impaired sleep-dependent gene expression in *Cacna1g* LoF animals.

Taken together, *Cacna1g* LoF alters gene expression patterns across light and dark periods, with a markedly greater number of DEGs and TI scores emerging during the light period, when animals are predominantly asleep. Moreover, suppression of ARG programs and disruption of key molecular pathways exhibit pronounced region- and period-specific patterns in *Cacna1g* LoF animals. Collectively, these findings underscore a critical role for *Cacna1g* in regulating brain activity and transcriptional programs during sleep across brain regions.

### *Cacna1g* LoF impairs diurnal dynamics of sleep-wake regulated genes

Sleep timing and duration are governed by two interacting biological processes: an intrinsic circadian oscillator and a homeostatic sleep-wake-driven process. Together, these processes coordinate gene expression differences between light and dark periods that are essential for normal brain function (*53*). Genes whose expression cycles independently of sleep pressure are regulated by intrinsic circadian rhythms and are referred to as intrinsic rhythmic genes (IRGs), such as *Bmal1* and *Cry1* (*53*). In contrast, genes sensitive to sleep deprivation are driven by the homeostatic sleep-wake cycle and are classified as sleep-wake genes (SWGs), such as *Homer1* (*54*) and *Paqr8* (*53*). To distinguish the effects of *Cacna1g* LoF on these two sleep-dependent regulatory processes, we examined the expression patterns of representative IRGs and SWGs. As expected, *Bmal1* and *Cry1* exhibited normal period-dependent expression across all genotypes, indicating preserved intrinsic circadian regulation (Fig. S7A). In contrast, *Homer1* showed the expected upregulation during the dark period in WT animals in the prefrontal cortex (*54*); however, this pattern was reversed in KO mice with aberrant upregulation during the light period. A similar disruption was observed for *Paqr8*, whose diurnal expression pattern was inverted across all four brain regions examined (Fig. S7B). Analyses from these genes suggest that *Cacna1g* LoF may impair diurnal gene expression pattern in SWGs.

To evaluate these effects at a systemic level, we analyzed gene sets exhibiting rhythmic expression across light and dark cycles (*50, 53, 54*), encompassing both SWGs and IRGs. We found that rhythmic gene expression programs were significantly disrupted by *Cacna1g* LoF, with the most pronounced effects observed in KO mice during the light period across all examined brain regions (**Fig. 4E**, table S3). Notably, genes typically upregulated during sleep (sleep-associated genes) were aberrantly enriched among the downregulated genes, whereas genes normally upregulated during wake (wake-associated genes) exhibited opposite enrichment (**Fig 4E**). Conversely, during the dark period, this pattern was reversed, exemplified by enrichments of wake-associated genes among downregulation and sleep-associated genes among upregulation in the prefrontal cortex and thalamus (**Fig. 4E**). We next distinguished between SWGs from IRGs (*53*), and found that *Cacna1g* LoF induced substantially greater perturbation in SWGs than in IRGs, with the strongest effects observed in the prefrontal cortex during both light and dark periods (**Fig. 4E**). These results indicate that *Cacna1g* LoF selectively disrupts sleep-wake homeostatic regulation rather than intrinsic circadian rhythmicity. The directionality of these changes, together with enrichment of GR- and SD-regulated genes during the sleep period, suggests that the brain adopts a sleep-like molecular signature. Importantly, this selective impairment of SWGs provides a mechanistic basis for the attenuated diurnal transcriptomic signature observed across all four brain regions in the *Cacna1g* KO mice.

We next sought to identify the upstream regulators of the sleep-wake gene (SWG) signal by performing transcription factor (TF) overrepresentation analysis (*64*) specifically on the leading-edge genes identified in GSEA (**Fig. 4E**, table S3). This approach allowed us to pinpoint the TFs with the greatest influence on the disrupted SWG landscape. *Atf6* and *Xbp1* were among the most overrepresented TFs driving wake-associated genes across all 4 brain regions, whereas *Pou4f1* was found to be overrepresented in the prefrontal cortex and hippocampus (**Fig. 4F**). No significant enrichment was observed for TFs regulating sleep-associated genes (data not shown). We next examined inferred TF activity (*65*). In WT and HT animals, *Atf6* activity (dark vs. light) was higher during the dark cycle; however, this pattern was significantly reversed in KO mice (**Fig. 4G**, table S4). A similar, albeit weaker and not statistically significant, trend was observed for *Xbp1* while changes in *Pou4f1* were comparatively modest. To determine whether these activity changes were reflected at the transcriptional level, we examined the expression of *Atf6* and *Xbp1* directly. *Atf6* expression remained stable across time periods, whereas *Xbp1* exhibited robust period-dependent changes across all four brain regions (Fig. S8A-C). Importantly, *Cacna1g* LoF significantly altered the diurnal dynamics of *Xbp1*, as evidenced by significant or near-significant genotype-period interactions in KO mice (Figure S8C).

This pattern, characterized by the altered expression of *Atf6* target genes in the absence of *Atf6* mRNA changes, alongside the direct upregulation of *Xbp1* expression, is highly consistent with the canonical activation of the Unfolded Protein Response (UPR). Specifically, it suggests that Atf6 protein is post-translationally activated via proteolytic cleavage to drive the transcriptional induction of *Xbp1* (*66, 67*). To test this, we next analyzed GO terms related to protein homeostasis and ER stress. During the light period, ER stress- and protein folding-related pathways were enriched among the upregulated genes in KO animals in the hippocampus, thalamus, and striatum (**Fig. 4H**). In the prefrontal cortex, enrichment patterns were more nuanced: among protein folding-pathways, only genes associated with the ER chaperone complex were significantly enriched among upregulated genes in the KO, whereas other protein-folding pathways were enriched among the downregulated genes in HT during the light period (**Fig. 4I**). To further examine directionality, we analyzed diurnal expression of these GO terms within each genotype. Consistent with the reduced inferred *Xbp1* transcriptional activity, the diurnal expression of genes in these pathways was generally not significantly enriched in WT or HT but was enriched in the downregulated genes in KO in the hippocampus, thalamus, and striatum (Fig. S8D). In contrast, the results in the prefrontal cortex were more mixed. Approximately half of the pathways, including those related to protein folding, protein maturation, and protein folding chaperones, were downregulated in both WT and KO but were not significant in HT. However, genes in pathways related to the ER chaperone complex and chaperone-mediated protein folding were specifically downregulated in KO but showed no changes in WT or HT (Fig. S8D). These findings highlight a region-specific dysregulation. Taken together, the dysregulation of genes transcriptionally regulated by *Atf6*/*Xbp1* could underlie the aberrant wake-associated gene expression in KO animals, particularly during the light period. These results suggest a potential link between transcriptional misregulation and cellular stress responses in *Cacna1g* KO mice.

### *Cacna1g* LoF induces robust transcriptomic changes in glial cells and dysregulates sleep-wake gene programs in excitatory neurons

To investigate the cell-type specific impact of *Cacna1g* LoF, we performed single-nucleus RNA sequencing (snRNA-seq) of the prefrontal cortex during the light period when bulk RNA-seq analyses revealed the most pronounced period-dependent transcriptomic alterations (Fig. S6A). Clustering analyses identified the expected major neural and glial cell populations, confirming accurate cell-type classification (see Methods, **Fig. 5A**, S9A upper). To assess genotype-dependent transcriptional changes within each cell type, we applied a pseudo-bulk differential expression approach, which enabled comparison across genotypes. The TRADE algorithm revealed a nominally significant genotype effect on transcriptomic-wide impact (**Fig. 5B**, table S5). Notably, the KO genotype produced a substantially greater global transcriptional impact than the HT genotype. Strikingly, among all cell types examined, glial cells, including astrocytes, microglia, and oligodendrocytes, exhibited the strongest transcriptomic-wide impact, as indicated by the highest impact scores. This finding was unexpected, given that *Cacna1g* expression is low or absent in these glial cells (Fig. S9A, lower), suggesting that *Cacna1g* LoF in neurons may elicit secondary, non-cell autonomous transcriptional responses in glial cells.

**Figure 5.**
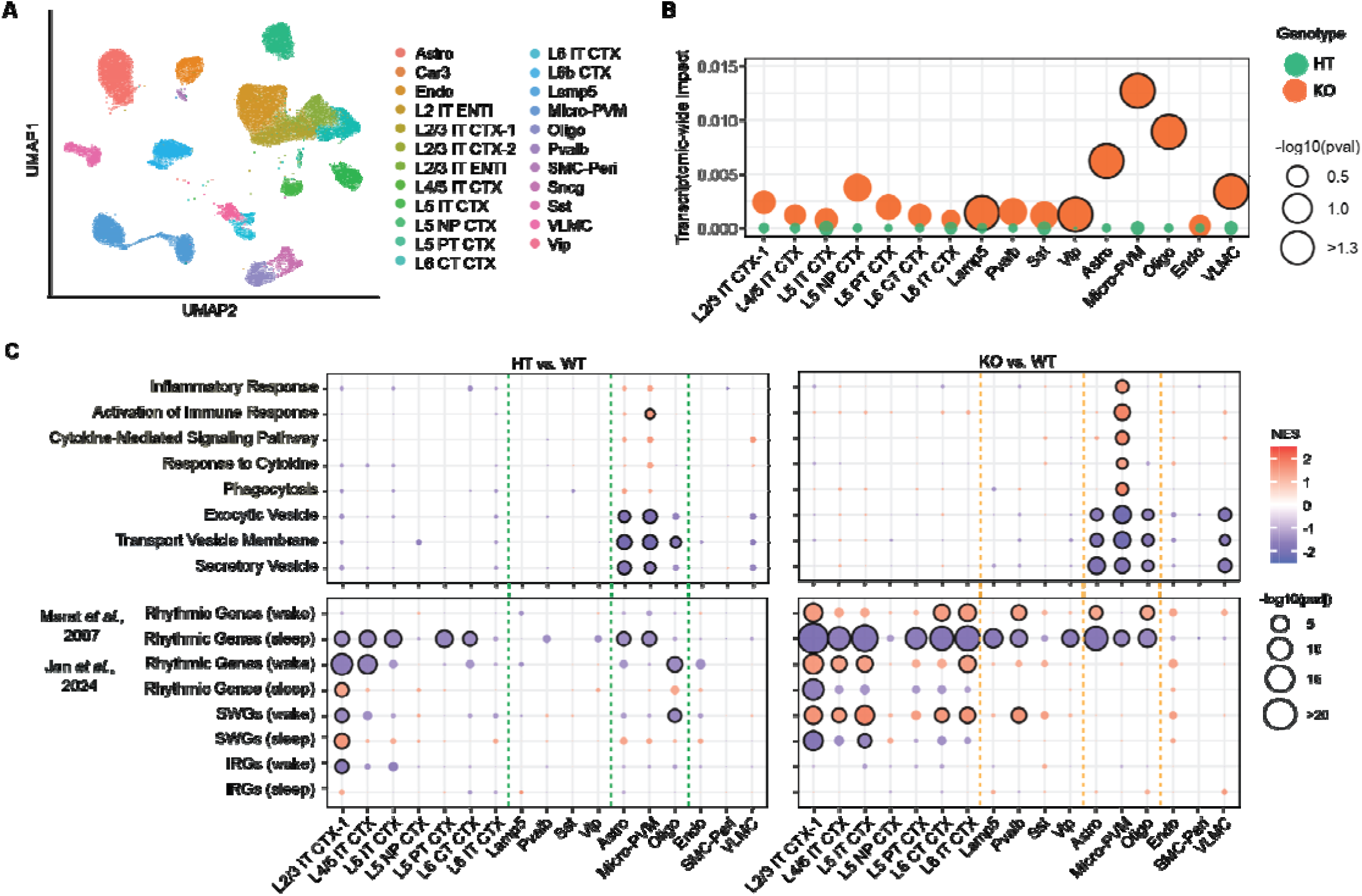
*Cacna1g* Loss of Function Impacts Transcriptomics in a Cell-Type Specific Manner. **(A)** UMAP visualization of cell types identified by snRNA-seq in the prefrontal cortex (PFC) of *Cacna1g* loss-of-function (LoF) mice and their WT littermates. **(B)** Transcriptome-wide impact of *Cacna1g* LoF across cell types. Black circle represents a given cell type with nominal significant (p<0.05) transcriptome-wide impact. **(C)** GSEA of gene ontology (GO) terms related to glial cell function and rhythmic gene sets in different cell types. The dotted lines separate major cell types: excitatory neurons (left), inhibitory neurons (center left), glial cells (center right), and others (right). CTX, cortex; L2–L6, layers 2–6; IT, intratelencephalic; NP, near-projecting; ET, extratelencephalic; CT, corticothalamic neurons; Lamp5, lysosomal-associated membrane protein 5 interneurons; Pvalb, parvalbumin interneurons; Sst, somatostatin interneurons; Vip, vasoactive intestinal peptide interneurons; Astro, astrocytes; Micro-PVM, microglia and perivascular macrophages; Oligo, oligodendrocyte; Endo, endothelial cells; SMC-Peri, vascular mural cells; VLMC, vascular and leptomeningeal cells.

To further elucidate cell-specific transcriptional changes, we performed GSEA using GO terms. In microglia from the KO animals, immune-related pathways, including inflammatory response and phagocytosis, which are strongly associated with microglial function, were significantly enriched among upregulated genes, consistent with an enhanced microglia activation (**Fig. 5C**, upper panel). Additionally, pathways related to exocytosis, including exocytic vesicle and secretory vesicle, were notably downregulated across all glial cell types, suggesting a broad disruption of glial secretory functions. Importantly, although these glial populations are more among the more abundant cell types in the dataset, other cell types with comparable abundances (Fig. S9A) did not exhibit enrichment of these pathways, supporting the specificity of these effects to glial cells (**Fig. 5C, upper panel**). Furthermore, GSEA on glucocorticoid receptor (GR)-regulated gene sets revealed upregulation of GR signaling pathways in both astrocyte and microglia in KO mice during the light period (Fig. S9B). Together, these findings indicate that *Cacna1g* KO mice induce pronounced and elective transcriptional remodeling in glial cells, which may contribute to the sleep-associated pathophysiology observed in *Cacna1g* KO mice.

We next examined the rhythmic gene expression associated with sleep-wake cycles across cell types. Consistent with our bulk RNA-seq results, wake-associated genes were enriched among upregulated genes across most of the cell types in KO animals, whereas sleep-associated genes were enriched among downregulated genes, indicating a blunting of normal sleep-wake dependent transcriptional regulation (**Fig. 5C, lower right panel**). These effects were particularly pronounced in L2/3 (layer 2/3) intratelencephalic (IT) excitatory neurons, which showed the strongest transcriptional changes, although this may be partially influenced by their relative abundance within the dataset (Fig. S9A). To determine whether wake-associated genes aberrantly activated in KO mice during the light cycle were also dysregulated in specific cell types and consistent across RNA-seq platforms, we intersected the GSEA leading-edge genes, the primary drivers of the “wake-state” signal in our bulk dataset (**Fig. 4E**, table S3), with those from snRNA-seq (**Fig. 5C**, table S5). Analysis revealed a set of overlapping genes between two data sets, such as *Shank2* and *Mamld1*, suggesting some overlap between the two methodologies (Fig. S9C). Notably, the involvement of IT neurons suggests that this neuronal subtype may drive the observed bulk transcriptomic changes associated with wake-state dysregulation in KO mice.

In summary, our snRNA-seq analysis reveals that *Cacna1g* LoF exerts a profound, genotype-dependent effect on the transcriptomic landscape of the prefrontal cortex, with distinct impacts on glial cell function, immune response pathways, and sleep-wake-dependent gene regulation. These findings underscores the complexity of *Cacna1g*’s role in brain function and identifies glial cells and excitatory neurons in the prefrontal cortex as key cellular substrates underlying the observed transcriptional disruption.

### *Cacna1g* LoF impairs molecular pathways related to protein homeostasis in synapse during sleep using proteomic analyses

Our transcriptomic analysis indicated that *Cacna1g* LoF impairs gene sets and pathways related to synaptic function (Fig. S6D), but also specifically highlighted a strong disruption in protein folding, chaperone activity, and the *Atf6/Xbp1*-driven UPR (**Fig. 4F-H**). This molecular signature of cellular stress predicts a subsequent collapse in the highly dynamic and vulnerable synaptic proteome (*68*). Additionally, the synaptic proteome is known to be sensitive to sleep-wake cycles (*3, 69, 70*). To further explore the impact of *Cacna1g* LoF at synapses, we performed a crude postsynaptic density (PSD) proteome analysis (*71*), focusing on the synaptic proteome of the HT and the WT animals across the light and dark periods in the prefrontal cortex and hippocampus. While the most pronounced UPR-related transcriptomic signatures were observed in KO mice, we focused our synaptic proteome analysis on the HT to identify the primary molecular vulnerabilities most relevant to the heterozygous state observed in human SCZ risk (*34*). We reasoned that while HT animals maintain enough homeostatic capacity to avoid the overt transcriptional ER-stress response seen in the KO (**Fig. 4**), the 50% loss of Ca_V_3.1 may still result in subtle, yet critical proteome remodeling at the synapse. Supporting this, we observed that the HT synaptic proteome was significantly disrupted despite the absence of a global transcriptomic collapse (**Fig. 6A**). This disruption was characterized by a distinct diurnal redistribution of synaptic protein expression across both the prefrontal cortex and hippocampus. Using RRHO2 to compare global synaptic protein rankings, we uncovered an inversion of protein enrichment in the prefrontal cortex. A large cohort of synaptic proteins that normally peaked during the dark period in WT animals showed significant enrichment during the light period in HT mice (**Fig. 6A**, upper). In contrast, minimal overlap was observed between WT and HT animals in the hippocampus, where diurnal protein patterns appeared largely decoupled (**Fig. 6A**, lower).

**Figure 6.**
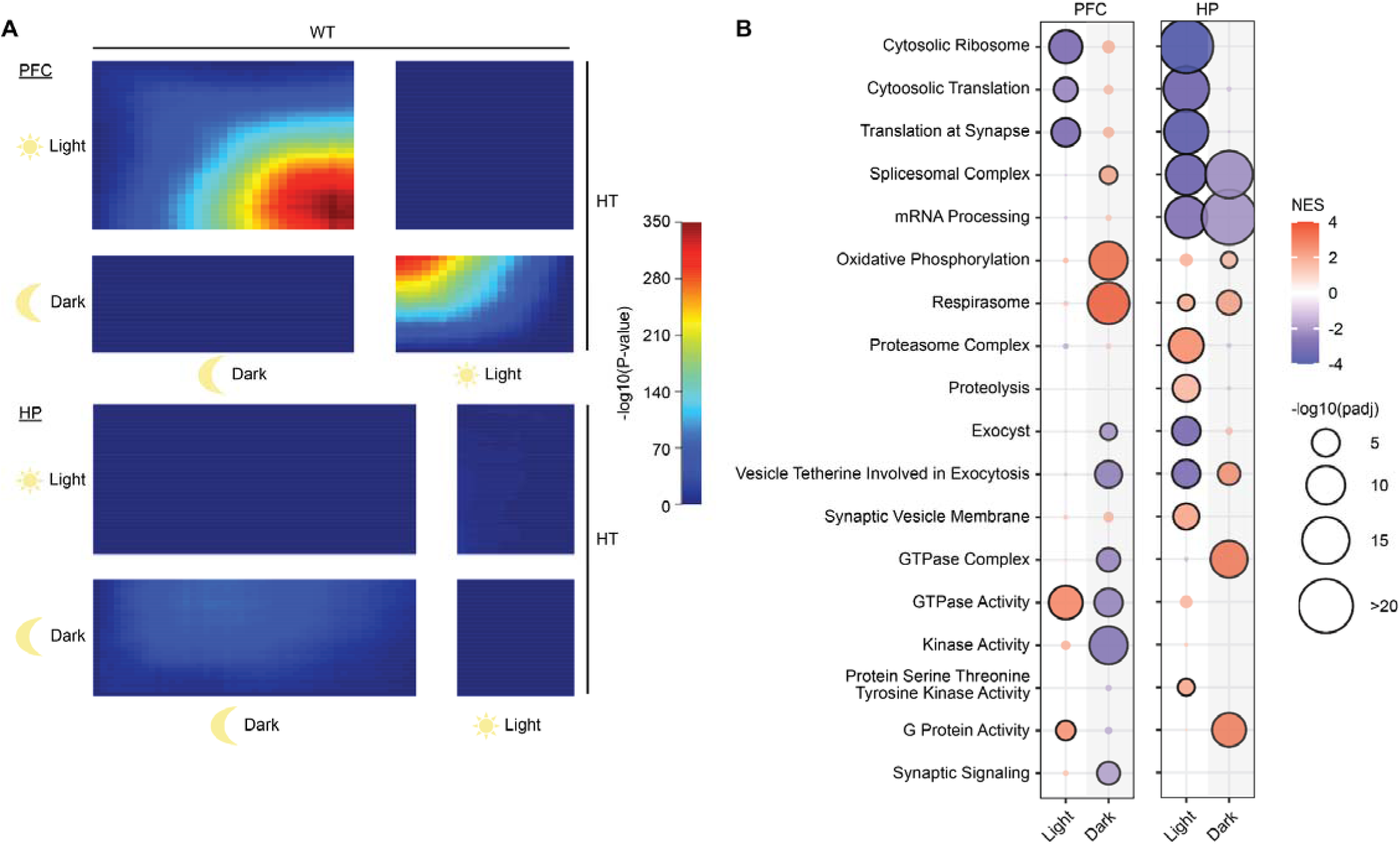
Cycle-Specific Impacts of *Cacna1g* Loss of Function on Molecular Pathways in the Prefrontal Cortex and Hippocampus Synaptic Proteome. **(A)** RRHO2 analysis comparing diurnal synaptic proteomic changes between WT and HT in the prefrontal cortex (PFC) and hippocampus (HP). **(B)** Differential expression of selected Gene Ontology (GO) terms for PFC and HP tissues between HT and WT animals in both light and dark periods.

Pathway analysis revealed that biological pathway changes occurred during both light and dark periods in both regions (**Fig. 6B**, table S7). Protein related to ribosome- and translation-related pathways were significantly enriched with reduced expression during the light period in both the prefrontal cortex and hippocampus, while mitochondria-related proteins, such as those related to oxidative phosphorylation, exhibited enrichment among the upregulated proteins during the dark period (**Fig. 6B**). In contrast, proteasome-related proteins were enriched among upregulated proteins during the light period in HT animals specifically within the hippocampus but had no significant change in the prefrontal cortex during the same period. Proteins associated with mRNA processing were consistently enriched among downregulation in the HT hippocampus across both light and dark periods, a pattern not observed in the prefrontal cortex (**Fig. 6B**). Synapse-related signal transduction proteins, including those involving GTPase activity, kinase activity, and G protein activity, were predominantly impaired during the dark period, showing enrichment among upregulated proteins in the hippocampus but among the downregulated proteins in the prefrontal cortex (**Fig. 6B**). Together, these findings demonstrate the central role of *Cacna1g*/Ca_V_3.1 in orchestrating widespread biological process changes at synapses across periods.

### *Cacna1g* LoF induces hyperactivity during dark cycle

Finally, we assessed whether *Cacna1g* LoF causes behavioral alterations using unsupervised behavioral profiling platforms. We first employed the home cage monitoring system to track animal activity in their home cage over 60 days, enabling us to examine the impact of *Cacna1g* LoF on circadian rhythms, as indicated by locomotor activity, in a more natural environment. Mice were group-housed after weaning and subsequently singly housed in the monitoring cage around postnatal day 28. Locomotion data from both HT and WT animals were recorded during the light and dark periods, revealing distinct activity patterns in the two phases, as expected. These findings suggest that the overall circadian rhythm remained intact (**Fig. 7A**). HT animals exhibited hyperlocomotion entering either the dark period (ZT12-14), or light period (ZT0-2), compared to WT littermates (**Fig. 7B**).

**Figure 7.**
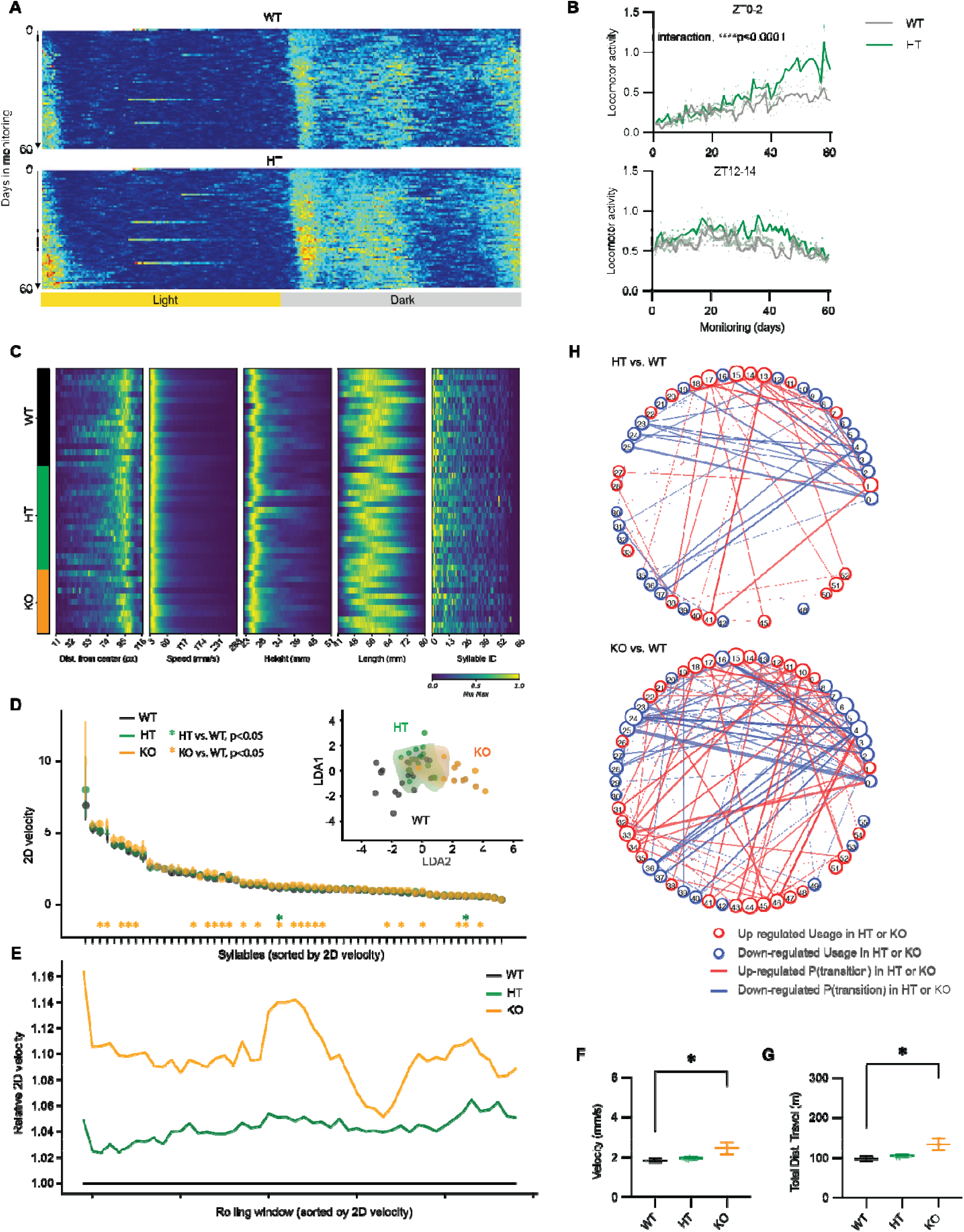
Unsupervised Behavioral Profiling Reveals Hyperactivity in *Cacna1g* Loss of Function Mice. **(A)** Heatmap showing locomotor activity of *Cacna1g* WT and HT mice in the home-cage monitoring system. Each bin represents the aggregated activity over a 5-minute interval. The y-axis spans 60 days of monitoring (postnatal day 28 to 98), with each row corresponding to a single day. Light and dark periods are indicated. **(B)** Average locomotor activity of *Cacna1g* WT and HT mice across 60 days, shown separately for light (upper) and dark (lower) periods. ****p < 0.0001, interaction analysis by two-way ANOVA. **(C)** Heatmap illustrating MoSeq position (defined as normalized distance from the arena center), speed, height, length, and syllable ID. Each row represents one session, comprising one mouse recorded on one experimental day within the arena. The heatmap is sorted by genotype, with each row normalized to the minimum/maximum value within the session. **(D)** Average syllable velocity in Cacna1g WT (*N*=17), HT (*N*=18), and KO (*N*=11) mice. Syllables are ordered by 2D velocity. ∗p < 0.05 by Kruskal-Wallis test. Data are mean ± 95% CI. Syllable labels were assigned by the analysis. Insert: LDA clustering of Cacna1g WT, HT, and KO mice using velocity-related parameters. **(E)** Relative average syllable velocity in Cacna1g HT and KO mice, normalized to WT. Syllables were sorted by mean velocity and averaged using a sliding window of 10 syllables. **(F, G)** Average velocity (F) and total distance traveled (G) in Cacna1g WT, HT, and KO animals. ∗p < 0.05 by Kruskal-Wallis test with Dunn’s multiple comparison correction **(H)** Transition graph depicting the alteration in syllable usage and transition probability in HT (upper) or KO (lower) mice, compared to WT mice.

Next, we employed motion sequencing (MoSeq), a machine learning-based behavioral profiling platform that divides animal behaviors into distinct syllables at a hundredth millisecond scale (*72*). We conducted MoSeq analysis during the dark period when animals are more active to determine whether the abnormal behaviors could be captured by an independent platform and gain a more detailed understanding of the alterations (**Fig. 7C**). MoSeq data revealed that, compared to WT littermates, *Cacna1g* HT and KO animals showed normal occupancy of the entire arena throughout the 30-minute recording period (Fig. S10A-B). There were no significant differences in the average distance to the center or height between genotypes (Fig. S10C-D), and these parameters did not cluster or separate genotypes (Fig. S10E-F), suggesting that *Cacna1g* LoF did not induce anxiety-like behavior in mice. However, when examining the usage of each syllable across different genotypes, we observed that *Cacna1g* LoF, particularly in KO animals, led to an increased usage of high-velocity syllables (Fig. S10G-H). Analysis of syllable velocities revealed that *Cacna1g* KO mice exhibited significantly higher velocities across nearly half of the syllables (**Fig. 7D-E**). Additionally, velocity-related parameters allowed for clustering of mice from the same genotype, with HT and KO animals distinct from WT (**Fig. 7D, inlet**). Furthermore, *Cacna1g* KO mice demonstrated significantly higher average velocity in the MoSeq arena, resulting in increased total distance traveled, indicative of hyperactivity (**Fig. 7F-G**).

By comparing syllable usage across all behaviors, we observed that certain behaviors, such as darting, were more prominent in HT and KO animals, as indicated by increased use of syllables 15 and 18 (see Methods, Fig. S10I). By contrast, behaviors related to turning, represented by syllable 23, were less frequent in HT and KO animals (Fig. S10I). Additionally, we detected significant changes in syllable transitions in both HT and KO animals, with these alterations following a gene-dose-dependent pattern. KO animals exhibited more pronounced and consistent directional changes compared to HT animals (**Fig. 7H**). Notably, the syllables showing enhanced transitions, such as 15 and 18, were also high-velocity syllables, which aligns with the observed hyperactivity in KO animals (**Fig. 7H**).

To conclude, two unsupervised behavioral profiling analyses revealed that *Cacna1g* LoF results in distinct alterations in motor activity and behavior, particularly in KO mice. While the circadian rhythm appeared largely intact, significant hyperactivity was observed in KO animals during active periods, alongside changes in specific movement patterns and behavior transitions, suggesting disruptions in spontaneous behaviors. These findings provide further insight into the functional consequences of *Cacna1g* LoF.

## DISCUSSION

The recent report suggesting that protein-truncating variants of *CACNA1G* contribute to an increased risk of SCZ motivated this study (*34*). Here, we define a translational molecular-to-network cascade linking rare genetic variation in *CACNA1G*, encoding the T-type calcium channel Ca_V_3.1, to core neurophysiological endophenotypes of SCZ. First, we establish the clinical relevance of *CACNA1G* by showing that individuals with SCZ who carry *CACNA1G* variants exhibit more severe sleep neurophysiological deficits, closely recapitulating the canonical SCZ versus control profile, yet are greater in magnitude (**Fig. 1**). These findings identify *CACNA1G* variation as a modifier of disease-relevant brain physiology in humans. Motivated by these human observations, we next leveraged a *Cacna1g* LoF mouse model to interrogate underlying mechanisms. Notably, *Cacna1g* LoF largely recapitulated the core EEG abnormalities observed in individuals with schizophrenia, including disruptions in spectral power, cortical coherence, sleep spindles, slow oscillations, and their coupling (**Fig. 2-3**), establishing strong cross-species convergence and validating this model as disease-relevant.

Despite largely preserved sleep macro-architecture, *Cacna1g* LoF profoundly disrupted diurnal regulation of brain activity and gene expression across the sleep-wake cycle. At the network level, loss of *Cacna1g* abolished normal 24-hour theta rhythmicity, indicating impaired temporal coordination of brain-state dynamics. At the molecular level, sleep-associated transcriptional programs were attenuated, while GR- and SD-regulated gene sets, which are typically associated with wakefulness and stress, were aberrantly enriched during the sleep period (**Fig. 4-5**). Together with disrupted sleep microstructure, these findings indicate that molecular programs normally entrained to wakefulness intrude into sleep, revealing a failure of sleep-dependent homeostatic regulation rather than intrinsic circadian timing. Importantly, this sleep-phase molecular dysregulation was accompanied by remodeling of the synaptic proteome, with widespread, period-dependent alterations in proteins involved in translation, proteostasis, and synaptic signaling (**Fig. 6**). This dysregulation persisted into wakefulness and manifested as a striking dissociation between brain state and behavior. During the active phase, wake-associated gene expression was blunted while sleep-associated gene programs remained inappropriately enriched, consistent with a “sleepy” or low-arousal molecular brain state (**Fig. 5**). In parallel, EEG measures revealed abnormal cortical power distributions, indicating altered network activation (**Fig. 2**). Paradoxically, despite this sleep-like neural and molecular signature, *Cacna1g* LoF animals exhibited behavioral hyperarousal, including increased home-cage activity and elevated high-velocity movement motifs detected by MoSeq (**Fig. 7**). Together, these findings identify *Cacna1g/*Ca_V_3.1 as a critical regulator of state-dependent brain activity across the sleep-wake cycle and demonstrate that its disruption drives a persistent mismatch between neural state and behavioral output, providing a mechanistic framework linking sleep dysregulation to circuit, synaptic, and behavioral abnormalities relevant to schizophrenia pathophysiology.

### *Cacna1g* is essential for coordinated microstructural and transcriptomic activity during sleep

*Cacna1g* LoF drastically reduces the spindle activities in NREM sleep (**Fig. 3B**). Spindle amplitude, duration and density are all significantly reduced in HT and KO animals, particularly in the frontal region. A reduction in sigma band power was previously reported in *Cacna1g*/Ca_V_3.1 LoF mice (*38*), consistent with our sigma band power analyses and event-based analyses. By contrast, (*37*) reported no spindle deficits in *Cacna1g* KO mice, which may stem from methodological differences in spindle detection algorithms or other factors. We employed wavelet-based, narrower frequency windows for analysis and utilized a more refined threshold detection method, leading to robust conclusions (*40*). Differences in recording locations may also contribute to variability in findings, as we do observe more robust finding in the frontal electrode compared to the parietal electrode (**Fig. 3B**, S3). In addition to sleep spindles, SO is the other cardinal oscillation during NREM sleep. *Cacna1g* LoF leads to an increase in both the number and duration of slow oscillation, which phenocopied impairment in patients of schizophrenia. Spindles arise from interactions between the thalamic reticular nucleus (TRN), and thalamocortical (TC) neurons during NREM sleep, while SO is thought to be generated in the layer 5 of cortical neurons (*73*). Ca_V_3.1 is expressed in layer 5 and 6 pyramidal neurons and at high levels in TC neurons (Fig. S9A, lower) and it could support spindle and SO oscillations directly during NREM sleep. Finally, we investigated the coupling between SOs and sleep spindles, crucial for memory consolidation and often impaired in SCZ (*74, 75*). Our result showed a decrease in coupling density and an increase in the coupled to uncoupled sleep spindles ratio, similar to what is observed in patients with SCZ. We also demonstrated that frontal region spindles couple with the down-sloping side of SOs, while parietal spindles couple with the up-sloping side. This spatial coupling mirrors findings from human studies (*8, 14*), where fast and slow spindles couple with the up and down slopes of slow oscillations, respectively. The regional difference in SO/Spindle coupling observed here suggests a similar spatial organization in mouse models, despite the lack of distinct spindle frequencies. Notably, the *Cacna1g* LoF leads to a shift in the coupling angle between spindles and SO, which is also observed in SCZ patients (*8, 14*). Our *Cacna1g* LoF model shows strong face validity for SCZ by phenocopying the major neurophysiological deficits of the disorder during sleep, as these observed key sleep deficits (altered SO, reduced spindles, impaired coupling) known to be robust SCZ biomarkers (AUC=0.93) (*8, 15*).

Our analyses introduced a temporal dimension by contrasting and comparing transcriptomic data from light and dark periods and examining sleep-wake dependent processes. We found that the LoF of *Cacna1g* significantly disrupted the expression of both activity-regulated genes (Fig. S6B, immediate early genes) and rhythmic genes (**Fig. 4F**), particularly those regulated by sleep and wakefulness, during the light cycle. The EEG analyses of sleep and wake states corroborate these findings, revealing that key oscillations, such as sleep spindles and slow oscillations, are severely disrupted in *Cacna1g* LoF animals during the same period. These cycle-specific disruptions highlight the critical role of *Cacna1g*/Ca_V_3.1 in shaping both the transcriptional and neurophysiological signatures during sleep. Ca_V_3.1 is known to drive synchronized oscillations in the thalamus during NREM sleep that then propagate to cortical regions (*76–78*). These oscillations are thought to promote activity-dependent transcription (*79*). In the absence of Ca_V_3.1, these oscillations are disrupted which leads to a reduction in the expression of activity-driven genes, which plays a pivotal role in synaptic plasticity, priming local synapses for strengthening or weakening (*80*). Genes related to synapses and synaptic function have been consistently implicated across both SCZ risk models and human post-mortem brain studies (*81–84*). Aligning with these findings, the reduced expression of IEGs during sleep in *Cacna1g* LoF animals suggests a disruption of normal sleep-dependent synaptic homeostasis. This conclusion is further supported by our synaptic proteomic pathway analysis, where it showed that pathways related to homeostasis, such as cytoplasmic translation and proteolysis, are enriched among downregulated proteins during the light period (**Fig. 6B**).

Further supporting a state of disrupted sleep microstructure, we identified heightened glucocorticoid (GC) signaling in *Cacna1g* KO mice, especially during the light period (Fig. S6F). This is consistent with the concurrent overrepresentation of sleep-deprivation-regulated genes and strongly suggests that *Cacna1g* LoF leads to hyperactive GC receptor (GR) signaling during sleep. This is particularly striking given that GR signaling normally follows a circadian pattern, peaking *before* wakefulness and troughing during sleep (*60*). Intriguingly, the cellular origin of this GC-related transcriptomic disruption appears distinct from that of the primary sleep-wake genes. Our PFC snRNA-seq analysis revealed that these GR-mediated gene sets were predominantly impacted in astrocytes (Fig. S9B), rather than in the excitatory or inhibitory neurons (like IT neurons) primarily affected by general sleep-wake dysregulation. Given that *Cacna1g*/Ca_V_3.1 itself shows low expression in glial cells relative to neurons (S9A, lower), this widespread transcriptomic impact on astrocytes likely represents a secondary consequence, potentially arising from altered neuro-glial interactions driven by the primary neuronal dysfunction. This interpretation resonates with human post-mortem studies implicating compromised neuro-glial communication in SCZ pathophysiology (*85*). Corroborating this notion of misregulated communication, pathway analysis highlighted enrichment in pathways related to exocytic vesicles and secretions among glia cells (**Fig. 6C**). The established links between GC signaling and theta waves (*86, 87*), which are themselves dysregulated in our model, highlight a complex interplay between neuroendocrine pathways, glial responses, network oscillations, and *Cacna1g*/Ca_V_3.1 function, underscoring its multifaceted contribution to disease biology.

### Loss of *Cacna1g* abolishes theta rhythmicity and impairs vigilant state transitions

Our analysis revealed that *Cacna1g* KO animals exhibit altered diurnal patterns of sleep-wake regulated genes (**Fig. 4E**). Interestingly, these genes were previously correlated with delta wave activity (*50, 53*), and we observed significant reduced delta power during NREM sleep in the KO animals compared to WT. The 24-hour rhythmicity of delta wave activity in the KO animals remained largely similar to that of WT (**Fig. 4A**), suggesting that this gene set is more sensitive to total delta power rather than the 24-hour rhythm of delta oscillations between wakefulness and sleep. Interestingly, the 24-hour rhythmicity of theta waves showed strong cosine patterns in WT and HT animals, while the KO animals displayed nearly constant theta power across 24 hours (**Fig. 4B**). This ablation of theta rhythmicity likely mirrors the altered expression of specific genes observed (**Fig. 4E**). These results suggest that although other factors like delta power may influence baseline gene expression, the 24-hour rhythmicity of theta power is a key driver of their temporal dynamics across the sleep-wake cycle.

Theta waves, which are prominent during both sleep and wakefulness, especially during REM sleep (*86, 87*), are known to mediate the expression of plasticity-related genes, potentially due to their higher frequency during sleep (*88*). In *Cacna1g* KO animals, we observed profound impairment of theta power in the frontal channel (**Fig. 2C**), suggesting the attenuation of sleep-induced increase or impairment of alertness during wakefulness. Specifically, we observed a loss of 24h rhythmicity in cortical theta power (**Fig. 4B**) and a trend toward reduced frontal-parietal theta coherence (**Fig. 2E**). Consistent with these network-level changes, our snRNA-seq analysis revealed that sleep-wake regulated genes were most significantly impacted within IT neurons (**Fig. 5C**), though this could be due to these neurons’ high abundance (Fig. S9A). IT neurons receive input from high order thalamic nuclei and modulatory subcortical regions, and project to other cortical areas (*89*). As such, this projection is known to integrate and distribute cortical information across widespread regions of the telencephalon (*90*). In KO animals, the aberrant expression of sleep-wake–associated genes during the light (sleep) phase suggests that key biological processes normally regulated during sleep are disrupted in these neurons. This disruption may impair both cell-intrinsic functions and coordinated activity across cortical networks. Supporting this interpretation, EEG recordings revealed reduced intracortical synchronization in KO animals (**Fig. 2D-F**), further indicating impaired cortical network dynamics that may stem from dysregulated IT neuron function during sleep.

Theta oscillations are also essential for organizing cortical and hippocampal networks during wakefulness, supporting processes such as attention, memory, and sensorimotor integration (*91*). The breakdown of 24-hour theta rhythmicity and coherence suggests that although *Cacna1g* KO mice are overtly awake and active (**Fig. 8G**), their underlying brain state, evidenced by altered neurophysiology and molecular signature, lacks the proper structured temporal coordination necessary for optimal wakefulness function. We propose that such persistent network dysrhythmia imposes a significant cellular burden, potentially inducing a chronic stress response. Our transcriptomic findings identified *Xbp1*, a key mediator of the unfolded protein response (UPR), as one likely driver of dysregulated sleep-wake genes, supporting this hypothesis. Established literature demonstrates that ER stress and UPR activation can severely disrupt synaptic plasticity and network function in key cognitive circuits (*92, 93*). Therefore, our finding of dysregulated *Xbp1* and an impaired diurnal rhythm of UPR pathways points towards a sustained cellular stress state in these mice. We therefore suggest a novel mechanistic link: *Cacna1g* LoF-induced network instability triggers or exacerbates cellular stress, which, in turn, further destabilizes brain vigilant state. Exploring this interplay between network oscillations and cellular homeostasis represents a critical avenue for understanding *Cacna1g*/Ca_V_3.1’s contribution to SCZ pathophysiology.

The impaired state transition also appears in our behavioral analysis. Both homecage monitoring and MoSeq analysis revealed behavioral alterations, notably hyperactivity, during the active dark cycle (**Fig. 8**), with these changes being most pronounced at the light-to-dark transition. MoSeq specifically identified shifts in high-velocity movements and their millisecond-scale transitions. Since sleep-related transcriptional and EEG disruptions often precede synaptic alterations that drive wakefulness behaviors, we propose that these behavioral abnormalities are consequences of the underlying molecular and neurophysiological deficits originating during sleep.

The impaired ability of *Cacna1g* LoF mice to stabilize sleep and wake states resonates with the pervasive sleep and circadian disruptions documented in SCZ. These are not mere comorbidities but are increasingly recognized as core pathophysiological features, often preceding psychosis onset and correlating with symptom severity (*94, 95*). The observed difficulties in our mouse model—manifesting as blunted 24-hour rhythms and destabilized brain states—mirror the human condition of unstable sleep-wake cycles, difficulties initiating/maintaining sleep, and reduced slow-wave sleep (*96*). This ‘state instability’ may extend beyond sleep and could represent a fundamental vulnerability underlying the cognitive inflexibility and, potentially, the fluctuating or abrupt nature of psychotic symptoms seen in SCZ (*97*). Therefore, the *Cacna1*g LoF model suggests a core deficit in the mechanisms maintaining stable brain states, potentially rooted in thalamocortical dysrhythmia, which bridges the gap between molecular dysfunction, sleep disturbances, and the broader symptomatology of SCZ.

### Conclusion, limitations, and future work

In conclusion, our study showed that *Cacna1g* LoF animals phenocopied the key neurophysiological deficits observed in individuals with SCZ, establishing this model as one with strong face validity for investigating SCZ-related pathophysiology. Integrated transcriptomics and synaptic proteomics analyses further reveal how *Cacna1g* modulates molecular, synaptic, and network-level activity during sleep. Together these results highlight the multifaceted role of *Cacna1g* in sleep-dependent processes that may contribute to SCZ pathophysiology. While our study provides important insights into the convergence of genetic regulation, neurophysiological dynamics, and sleep, we did not directly assess the impact of *Cacna1g* LoF on sleep-dependent memory consolidation, or other cognitive behaviors. Nevertheless, extensive prior work has addressed the correlational and sometimes causal links between sleep spindles, slow oscillations, and cognition. Future studies will build on this foundation to examine how *Cacna1g*-mediated disruption in sleep architecture and thalamocortical dynamics influence aspects of cognitive and behavioral output, thereby advancing our understanding how genetic perturbations of sleep-regulatory circuits contribute to the risk and development of psychiatric disorders.

## Supplementary Figures and Legends

**Figure S1.**
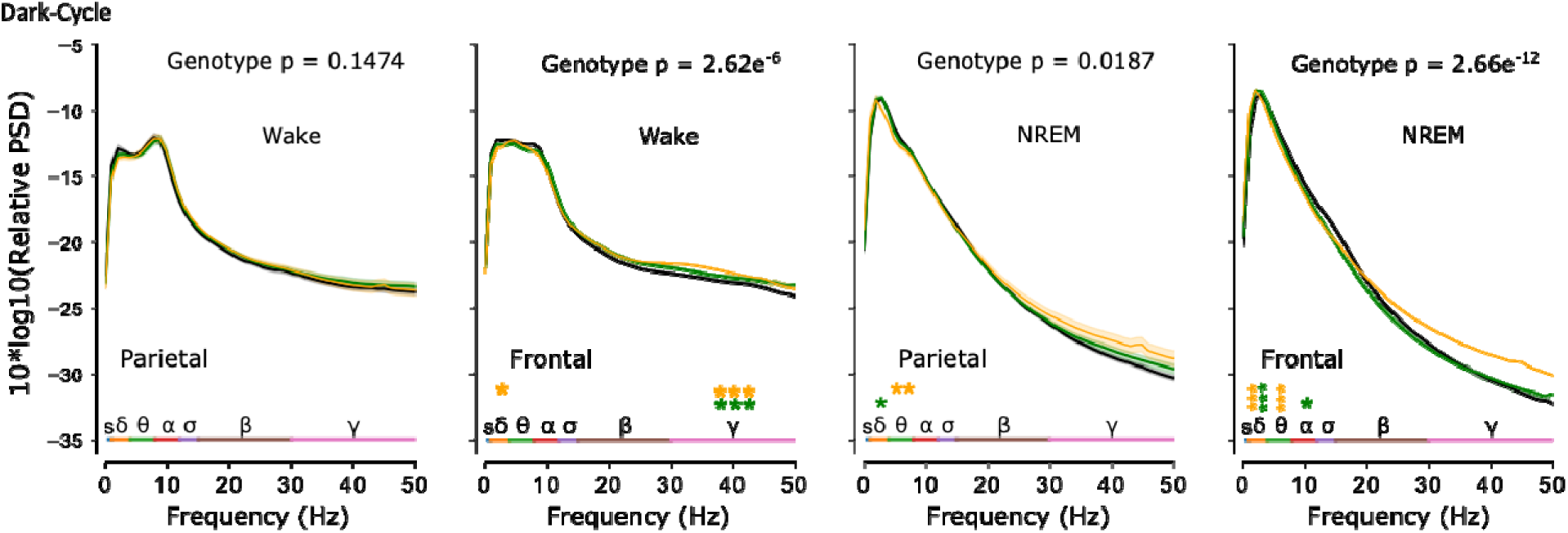
Related to Figure 2. Power spectrum comparisons during wakefulness and NREM sleep in parietal and frontal regions during the dark period. Thick lines indicate the mean, with shaded areas representing the standard error. Orange and green asterisks denote significant differences in specific frequency bands between KO and WT, and HT and WT animals, respectively. Statistical significance: *p* < 0.05 (linear mixed model followed by Tukey’s multiple comparisons test).

**Figure S2.**
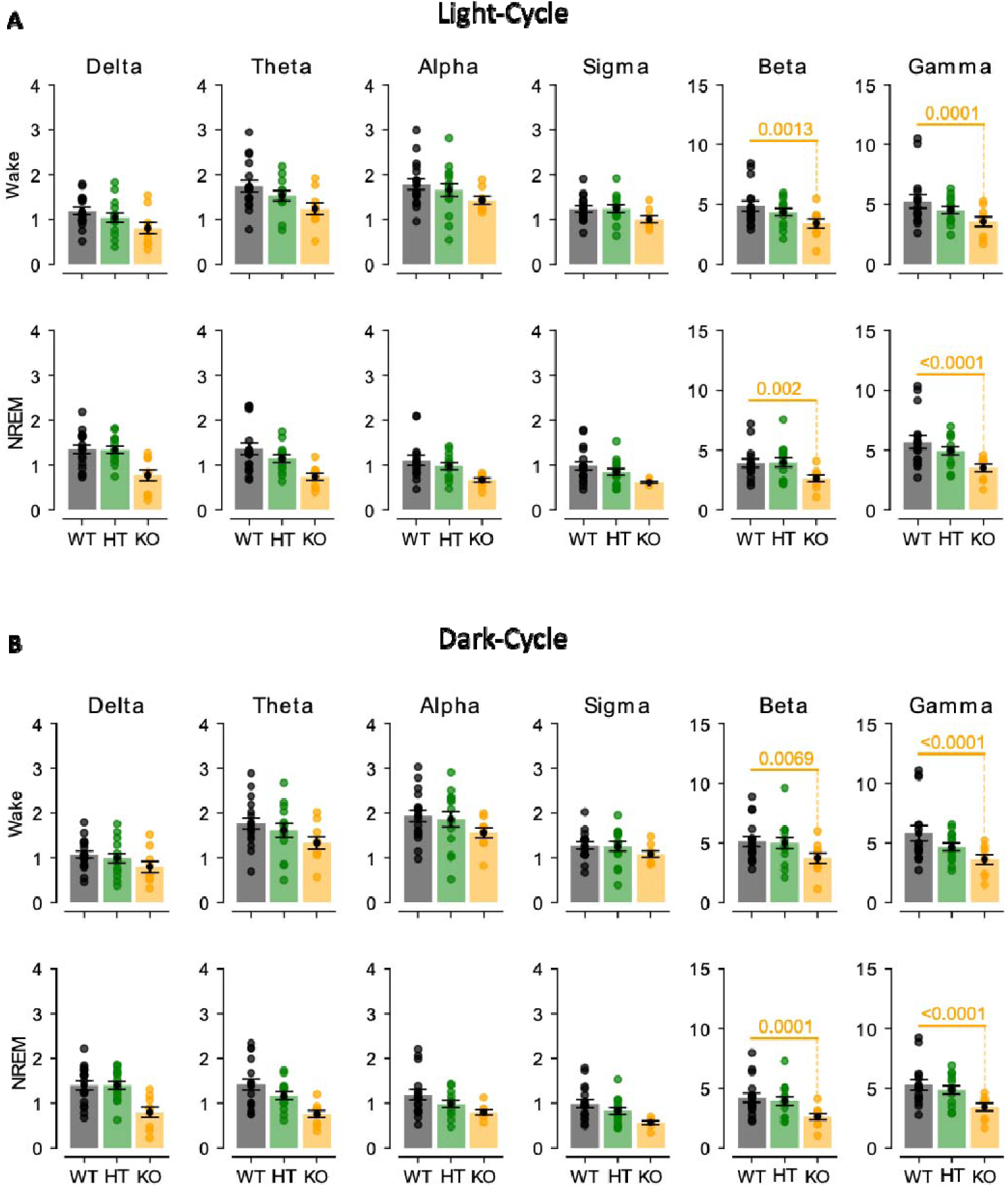
Related to Figure 2. Coherence comparisons between groups across frequency bands during the light (A) and dark (B) periods. Thick lines represent group means, with shaded areas indicating the standard error. Orange and green p-values denote significant pairwise differences between WT and KO, and WT and HT groups, respectively.

**Figure S3.**
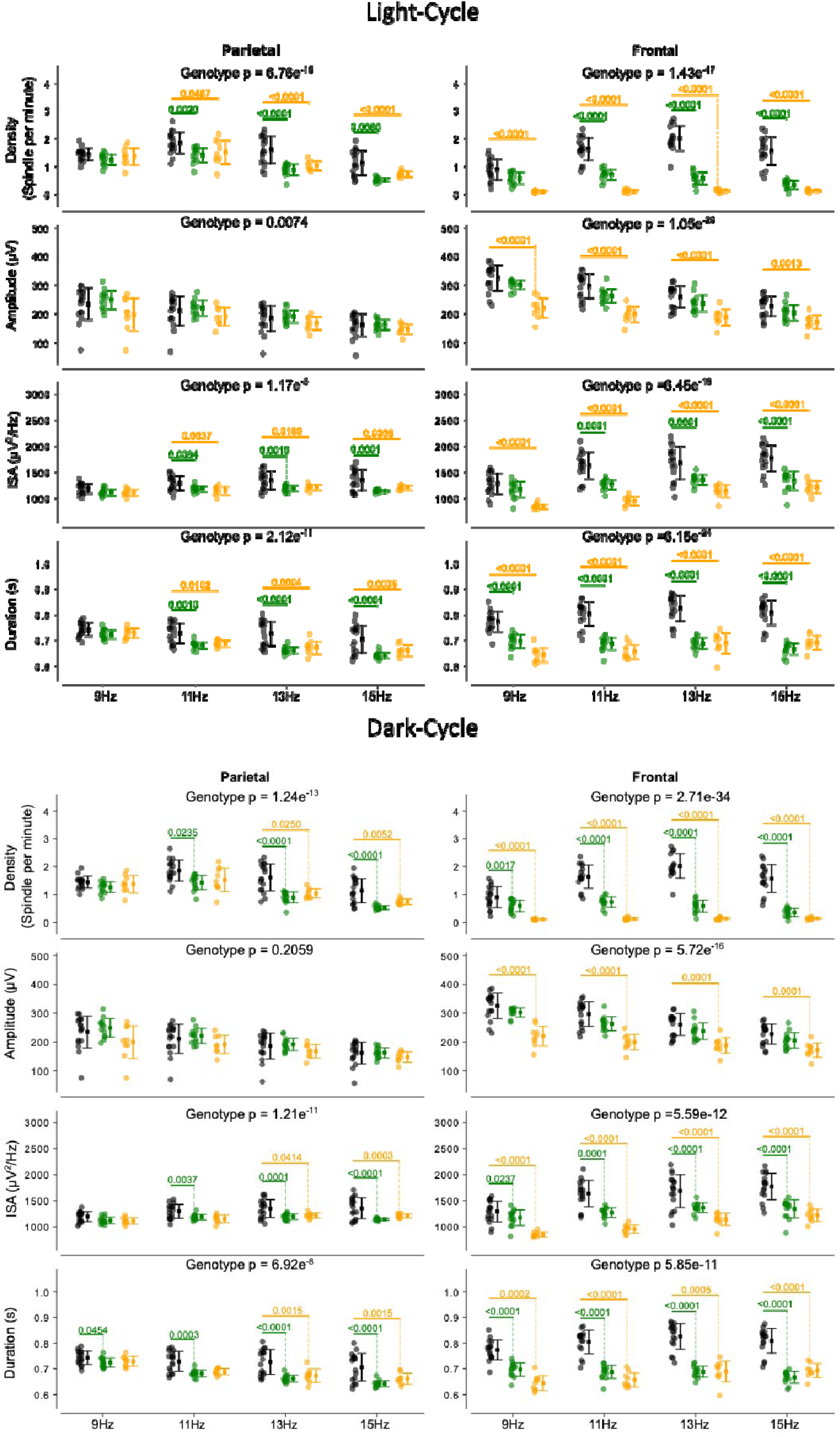
Related to Figure 3. Comparison of sleep spindle parameters (density, amplitude, integrated spindle activity, and duration) across genotype groups at 9, 11, 13, and 15 Hz during the light (A) and dark (B) periods.

**Figure S4.**
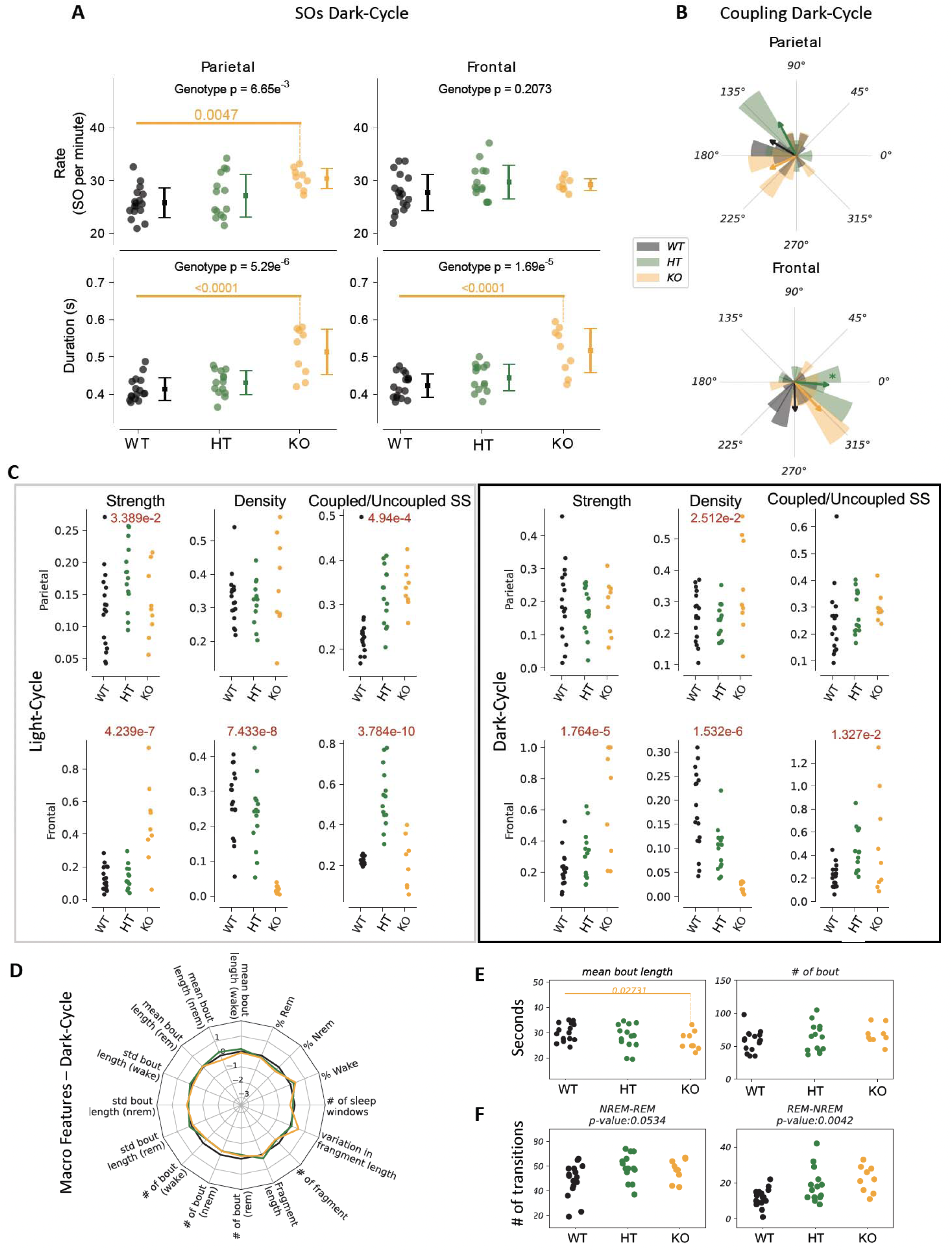
Related to Figure 3. **(A)** Comparison of slow oscillation (SO) rate and duration across groups during the dark cycle. Mean ± standard deviation is shown. **(B)** Group comparisons of spindle-SO coupling based on circular means during the dark cycle. **(C)** Comparison of coupling strength, spindle density, and coupled-to-uncoupled spindle ratio in parietal and frontal channels during both light and dark cycles. **(D)** Z-scored comparison of macro sleep features during the dark cycle across genotype groups. **(E)** Group comparisons of NREM bouts shorter than 60 seconds during the light cycle. **(F)** Number of transitions between NREM-REM and REM-NREM across genotype groups during the light cycle.

**Figure S5.**
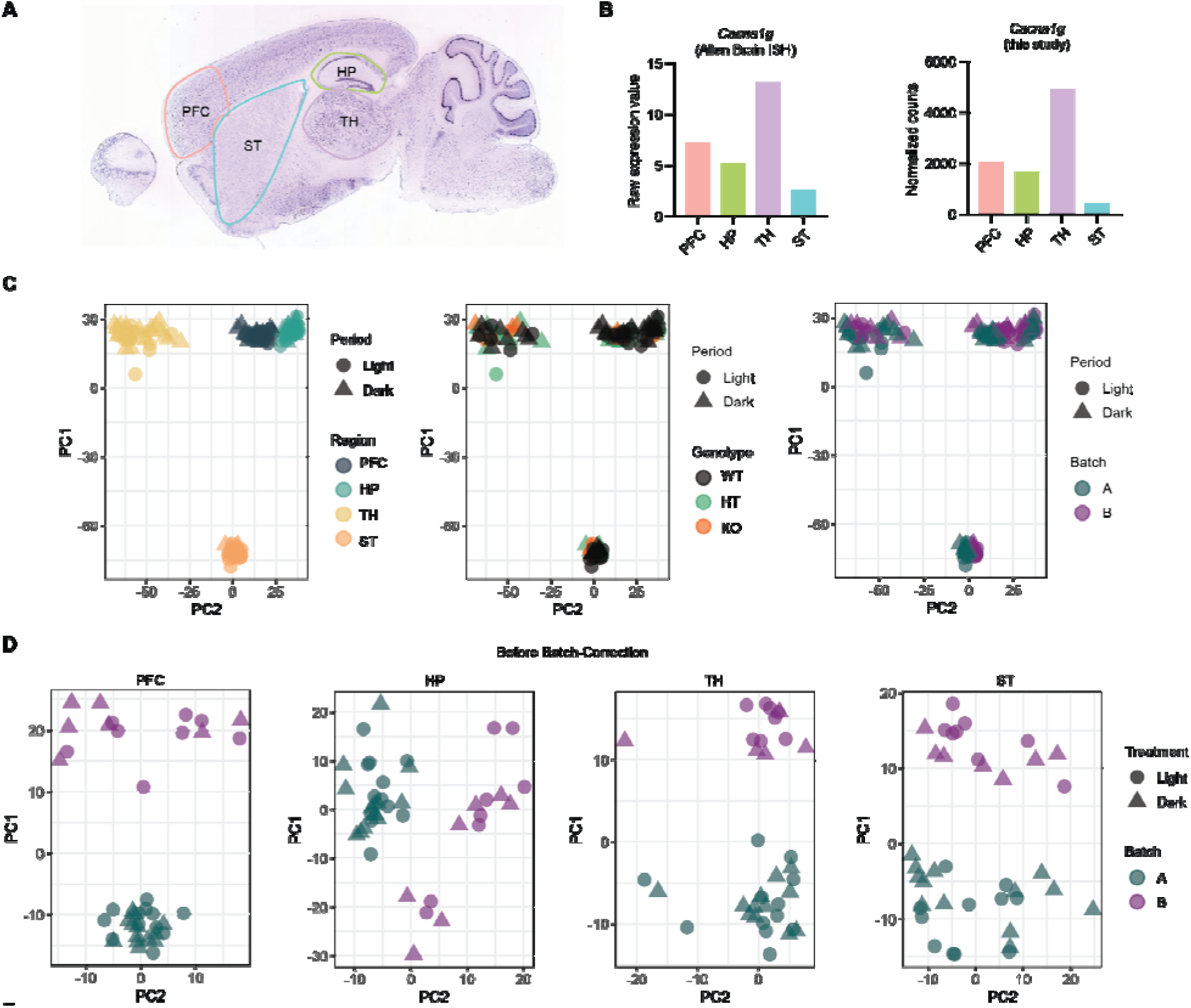
Related to Figure 4. **(A)** In situ hybridization image of *Cacna1g* from the Allen Brain Atlas (https://mouse.brain-map.org/gene/show/12076). **(B)** Raw expression values of *Cacna1g* from the Allen Brain Atlas (left) and normalized counts from RNA-seq in this study (right). **(C)** Principal component analysis (PCA) of individual samples across brain regions, with clustering by brain regions (left), genotypes (middle), and batches (right). **(D)** Principal component analysis (PCA) of individual samples before batch correction.

**Figure S6.**
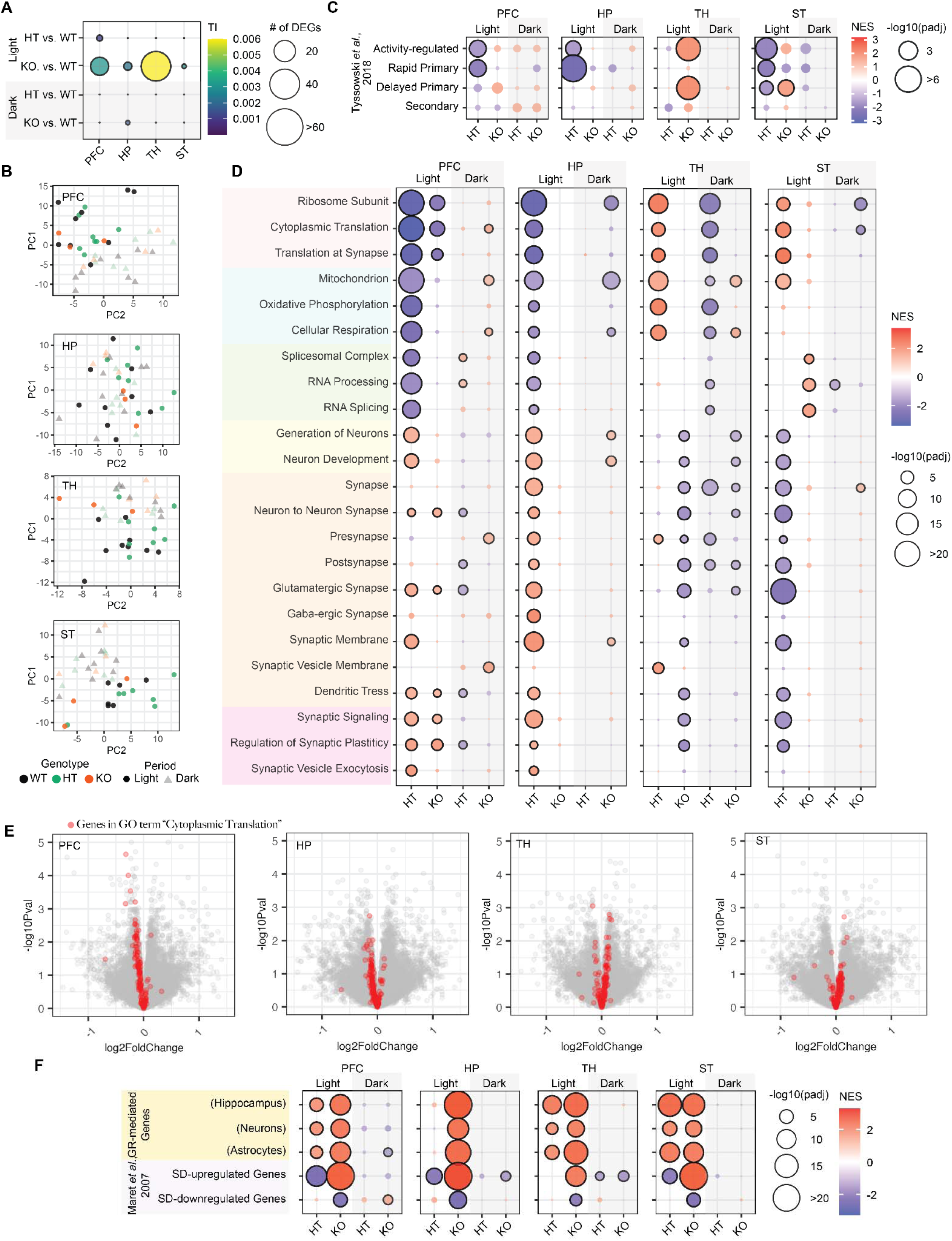
Related to Figure 4. **(A)** Number of differentially expressed genes (DEGs) and transcriptome-wide impact (TI) in *Cacna1g* LoF mice compared to WT controls across brain regions and diurnal phases. **(B)** Gene set enrichment analysis (GSEA) of activity-regulated gene sets in *Cacna1g* HT and KO mice compared to WT within each light-dark cycle. HT: HT vs. WT; KO: KO vs. WT. **(C)** Principal component analysis (PCA) of bulk RNA-seq profiles from PFC, HP, TH, and ST in *Cacna1g* LoF mice during light and dark phases. **(D)** Gene set enrichment analysis (GSEA) of Gene Ontology (GO) selected terms in HT and KO mice relative to WT during light and dark phases. HT: HT vs. WT; KO: KO vs. WT. **(E)** GSEA of gene sets associated with glucocorticoid signaling and sleep deprivation in HT and KO mice compared to WT.

**Figure S7.**
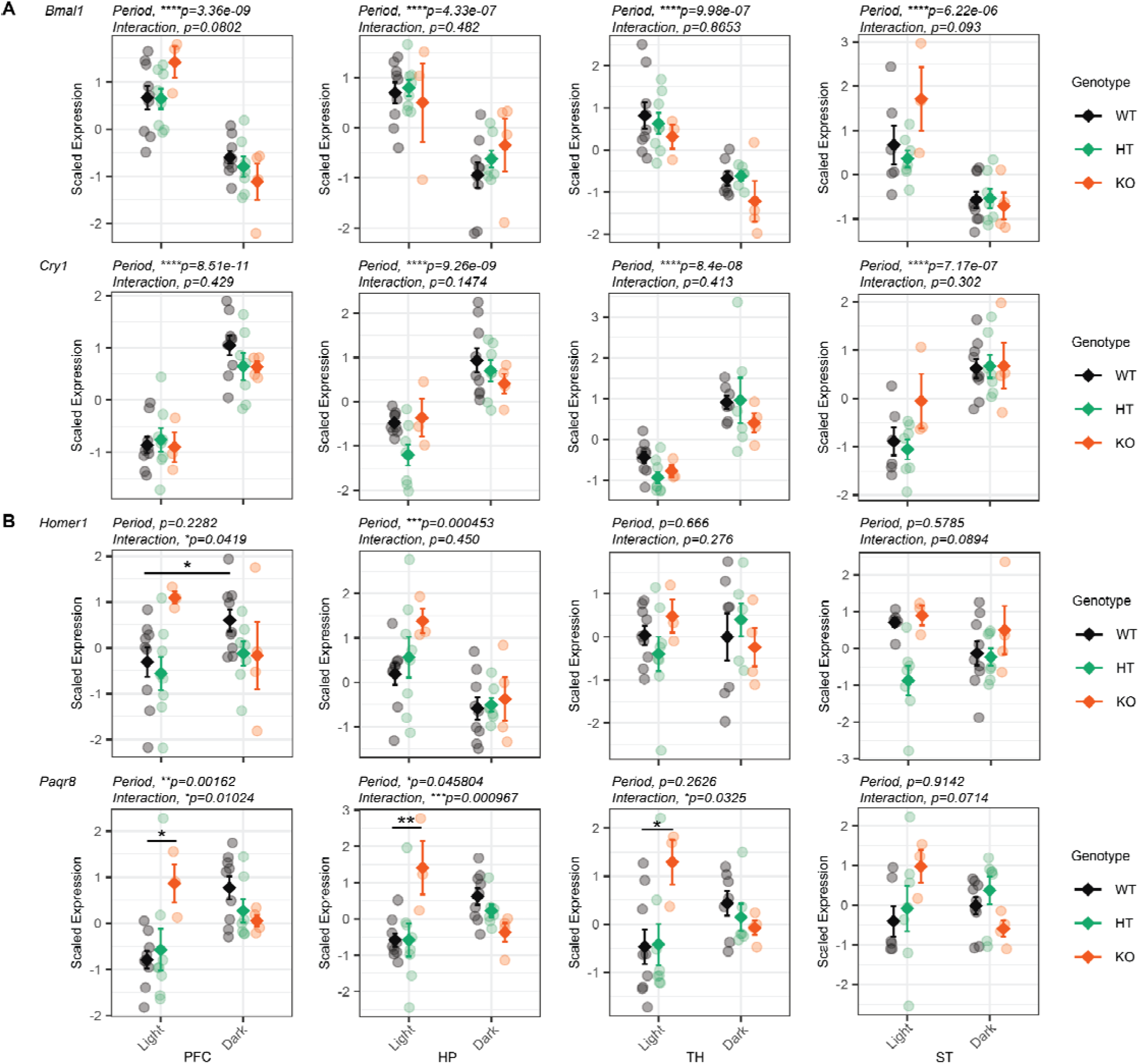
Related to Figure 4. **(A–B)** Scaled expression of core circadian genes *Bmal1* and *Cry1* (A), and sleep/wake-associated genes *Homer1* and *Paqr8* (B) across light-dark cycles in *Cacna1g* WT, HT, and KO mice in four brain regions. Significance was determined using a two-way ANOVA with Genotype (WT, HT, KO) and Period (Light, Dark) as main factors. The Period p-value assesses diurnal difference across all mice. The Interaction p-value specifically tests whether the effect of *Cacna1g* LoF is dependent on the diurnal phase. All reported ANOVA p-values are nominal. If a significant interaction was detected, post hoc pairwise comparisons were performed to identify specific genotype differences within each period or period difference within genotype. These post hoc tests were conducted using the emmeans package in R, with p-values adjusted for multiple comparisons using the Bonferroni correction. *p<0.05, **p<0.01, ***p<0.001, and ****p<0.0001.

**Figure S8.**
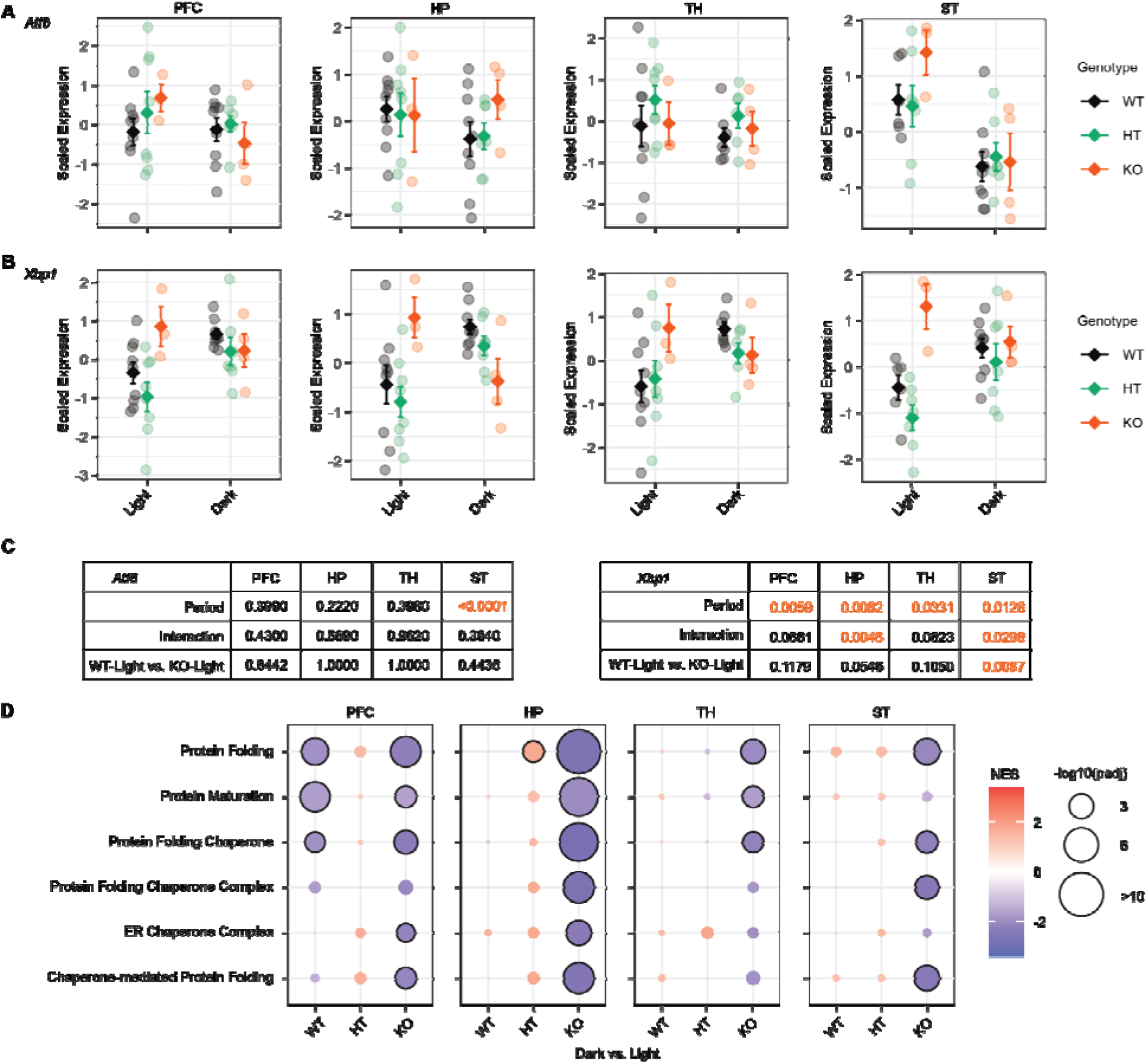
Related to Figure 4. **(A–B)** Scaled expression of transcription factors *Atf6* (A) and *Xbp1* (B) across light-dark cycles in *Cacna1g* WT, HT, and KO mice in four brain regions. **(C)** Two-way ANOVA summary for *Atf6* and *Xbp1* expression across genotype and cycle; comparisons with *p*lJ<lJ0.05 are highlighted. **(D)** GSEA of diurnal expression changes in genes involved in protein folding and maturation pathways, performed separately within each genotype (WT, HT, KO; dark vs. light).

**Figure S9.**
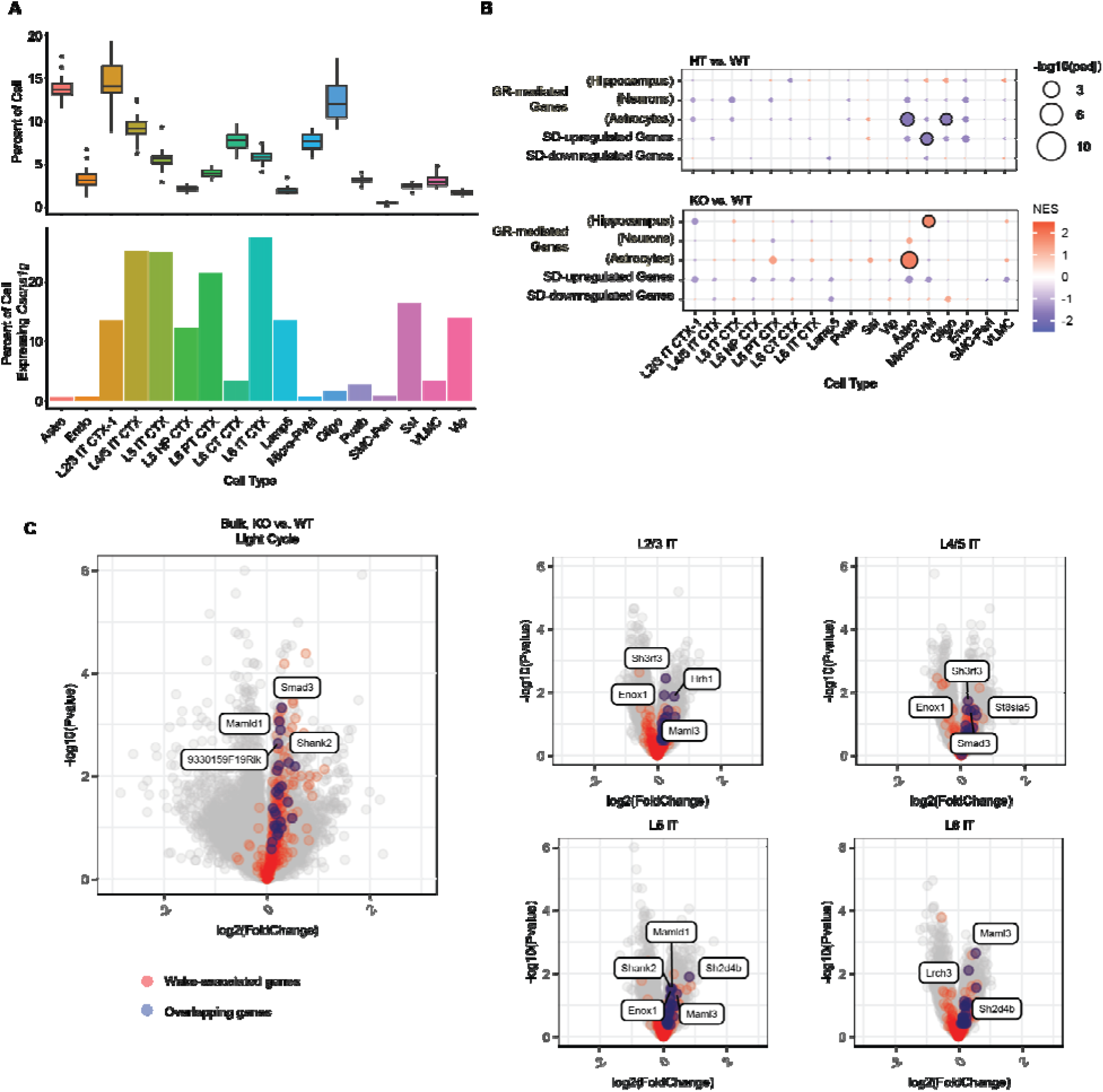
Related to Figure 5. **(A)** Proportional representation (upper) and *Cacna1g*-expressing cells (lower) of each major cell type across all samples. **(B)** GSEA of gene sets regulated by glucocorticoid receptor (GR) signaling and sleep deprivation (SD) in HT and KO mice compared to WT. **(C)** Volcano plots highlighting overlapping wake-associated genes (in red) identified in bulk RNA-seq (KO vs. WT during the light cycle) and in transcriptomic profiles of excitatory IT neurons from layers 2/3 (L2/3 IT), 4/5 (L4/5 IT), 5 (L5 IT), and 6 (L6 IT). Overlapping genes across bulk and sn RNA-seq were highlighted in blue.

**Figure S10.**
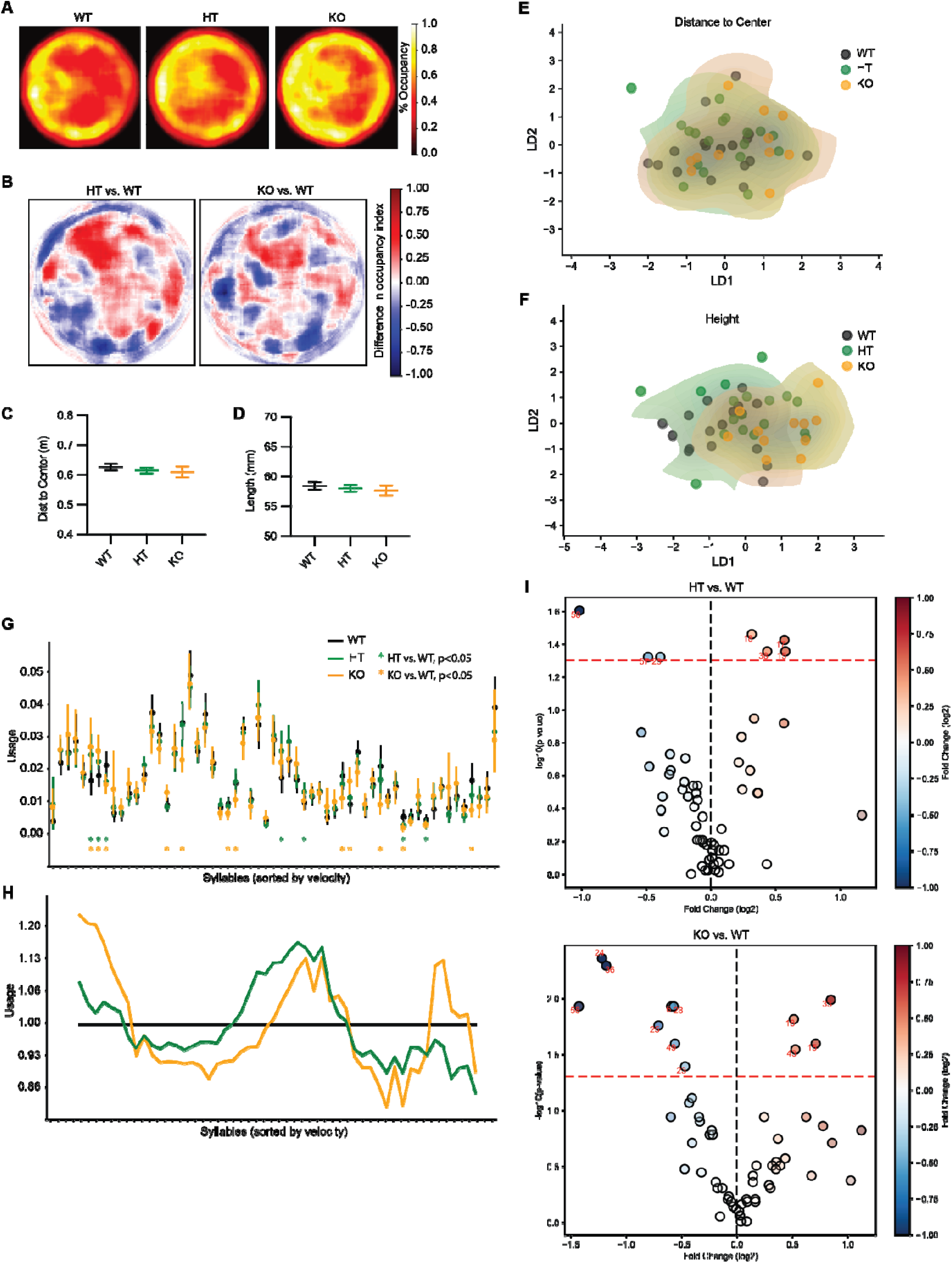
Related to Figure 7. **(A)** Heatmap showing occupancy percentages across the MoSeq recording period for Cacna1g WT, HT, and KO mice. **(B)** Differential occupancy heatmaps comparing HT vs. WT and KO vs. WT. **(C-D)** Average distance to the center (C) and average body length (D) across genotypes. **(E-F)** Linear discriminant analysis (LDA) clustering of WT, HT, and KO mice using features related to distance to the center (E) and body length (F). **(G)** Average syllable usage across genotypes, sorted by 2D velocity. *p* < 0.05 (Kruskal–Wallis test). Data are presented as mean ± 95% confidence interval (CI). **(H)** Relative syllable usage in HT and KO mice, normalized to WT. Syllables are ordered by mean velocity and averaged using a sliding window of 10 syllables. **(I)** Volcano plot showing differential syllable usage in HT vs. WT and KO vs. WT. Color indicates log2 fold change; red dotted line marks the *p* < 0.05 threshold.

## Supplementary Tables

Table S1 Differentially expressed genes on *Cacna1g* LoF bulk RNA-seq

Table S2 GSEA of *Cacna1g* LoF bulk RNA-seq

Table S3 Curated gene sets and GSEA of *Cacna1g* LoF bulk and sn RNA-seq

Table S4 decoupleR analysis inferring transcription factor activity

Table S5 Differentially expressed genes in *Cacna1g* LoF sn RNA-seq

Table S6 GSEA of *Cacna1g* LoF sn RNA-seq

Table S7 GSEA of *Cacna1g* LoF synaptic proteomics

## Materials Availability

Requests for materials and further information should be directed to the lead contact.

## Data and Code Availability

- Raw bulk RNA sequencing FASTQ files are deposited in the NCBI Gene Expression Omnibus (GEO) database (GSE299731). All datasets will be made publicly available upon acceptance of the manuscript.
- This study does not report any original code. The code for analysis of EEG, bulk RNA-seq and Moseq is in GitHub (https://github.com/ahmetassan25/Asan_et_al_2025).
- Any additional information required to reanalyze the data reported in this paper is available from the lead contact upon request.

## Acknowledgements

This research was funded by the Stanley Center for Psychiatric Research (J.Q.P. and J.Z.L.). The authors thank Dr. Richard P. Schiavoni (The Biopolymers and Proteomics Facility, Koch Institute For Integrative Cancer Research at MIT) for his assistance with proteomic experiments; Dr. Jixiang Zhang for his assistance with snRNA-seq experiments; Ms. Sahana Natarajan, Deeksha Misri, and Lakshanyaa Kannan for their support in MoSeq; and Dr. Bryan J. Song for his help with homecage monitoring. The authors also appreciate Dr. Kris Dickson for her feedback and suggestions during manuscript preparation, as well as all members of the Pan lab for their valuable support and contributions.

## Author Contributions

R.L. designed, conceived and analyzed the bulk, synaptic proteomics, and behavior studies. R.L. and A.S.A. analyzed the behavior studies. S.K. and H.H analyzed the sequencing from the GRINS cohort and identified the individual carrying CACNA1G variants. N.K. and S.M.P. analyzed the GRINS EEG data. N.G., E.Y., and S.C. conducted the mouse EEG recording and A.S.A. analyzed the EEG data. N.H., prepared the snRNA-seq libraries with supervision from J.Z.L., and S.K.S, and J.Z.L conducted the analysis. T.Y. and T.T.Z conducted the behavior experiments. L.Y., established the *Cacna1g* LoF mouse colony. R.L. and J.Q.P. conceptualized the study, and A.S.A., R.L., and J.Q.P. wrote the manuscript. All authors read and approved the final manuscript. This work was supported by the Stanley Center for Psychiatric Research.

## Methods

### Sequencing and variant calling, and quality control for GRINS samples

For the GRINS cohort, we generated sequencing data using the Blended Genome Exome (98) and DRAGEN protocol with PCR-free whole exome sequencing (WES) (99). Exome libraries were prepared using the Twist Alliance Clinical Research Exome kit, and whole-genome sequencing (WGS) and WES libraries were blended at a ratio of 67% WGS to 33% WES. The final blended pools were sequenced on an Illumina NovaSeq X. Joint calling of samples was performed using the Illumina DRAGEN pipeline². Variants were initially called per sample in gVCF mode, and sample-level gVCFs were then merged and jointly genotyped using the DRAGEN joint-calling workflow. For variant-level quality control, we started from the Genomic Variant Store (GVS) callset and split multiallelic sites into biallelic variants, retaining only sites with a PASS status in the Variant Extract-Train-Score (VETS) model. We removed variants located in low-complexity regions, telomeres/centromeres, and segmental duplication regions, and restricted analyses to the TWIST exome target intervals. We then applied filters to retain variants with call rate ≥ 0.95 and to remove monoallelic (invariant) sites. For genotype-level quality control, we excluded calls with genotype quality (GQ) < 20 or read depth (DP) < 10. Among heterozygous genotypes, calls with allele balance (AB) < 0.2 or AB > 0.8 were removed, and among homozygous alternate genotypes, calls with AB < 0.8 were excluded.

### Animals

Animal experiments adhered to Broad Institute IACUC guidelines and the NIH Guide for the Care and Use of Laboratory Animals. *Cacna1g* loss-of-function (LoF) mice, generously provided by Dr. Slobodan Todorovic (University of Colorado School of Medicine, Anschutz medical campus) and originally described by Petrenko et al (*100*), were maintained on a C57BL/6J background. Wild-type (+/+), heterozygous (+/-), and knockout (-/-) littermates for experiments were generated by crossing heterozygous mice. Mice were housed in AAALAC-approved facilities with a 12-hour light/dark cycle and ad libitum access to food and water. The first cohort of mice was generated for bulk RNA-seq and synaptic transcriptomics. Brain tissue samples were collected at the age of 10-14 weeks, during 2h entering either light or dark cycle, ZT 2-4 and ZT 14-16, respectively. The prefrontal cortex and hippocampus were dissected, one hemisphere for bulk RNA sequencing and the other for synaptic proteomics. Striatum and thalamus were dissected as well, combining both hemispheres to ensure sufficient RNA for bulk RNA sequencing. A second cohort of mice was generated for snRNA-seq and the tissues were harvested at the age of 6 weeks during light cycle, ZT 2-4. Only male mice were used in this study.

### EEG/EMG Implant Surgery

The animals were initially anesthetized using 4% isoflurane in a sedation chamber, with anesthesia levels subsequently maintained between 1% and 2% during the surgical procedure as previously described (*40*). Anesthesia depth was continually monitored by assessing the pinch reflex. Following the initial anesthesia, each animal was placed in a stereotaxic frame with their head fixed. Body temperature was maintained at 38 degrees throughout the procedure. The hair over the top of the head was removed and skin was cleaned with alcohol and antiseptic solution. A midline incision was made and connective tissue over the skull is excised. Given that spindle activity in the frontal and parietal regions exhibits distinct frequency characteristics and serves different functions in humans (*14*), electrodes were positioned in these regions to investigate whether similar characteristics are present in our study. Two shallow holes were opened at the coordinates (+1.5 AP, 1.5 ML to Bregma / −1.5 AP, 2.0 ML to Bregma) using a drill, and EEG screw electrodes (Pinnacle 8403) were placed into these holes. Two extra holes were made at the posterior part of the skull (bilaterally −1 AP, 2 ML to Lambda) over the cerebellum to insert the reference and ground electrodes. The screws were connected to a headmount (8201-SS Pinnacle) to allow for attachment to the recording system. For monitoring muscle activity, EMG wire electrodes were bilaterally implanted in the neck (nuchal) muscles. Connectors were fixed to the skull using dental cement. Following the surgery, animals were given a recovery period of 2 weeks. A total of forty mice (17 *Cacna1g+/+* (WT), 14 *Cacna1g+/-* (HT), and 9 *Cacna1g-/-* (KO)) had EEG implantation.

### EEG Recording

Once the animals had fully recovered from the surgery, EEG and EMG activity were recorded using a tethered system (Pinnacle 8200-K1-SL) connected to the head-mounted connectors. Recordings were conducted from freely moving animals. The recording sessions spanned a total of 24 hours, consisting of a 12-hour light-cycle and a 12-hour dark-cycle after 24 hours of habituation in the recording chamber. The recording system had a sampling rate of 1000□Hz and applied signal filtering (0.5–100□Hz bandpass for EEG; 10–1□kHz bandpass for EMG) (Pinnacle Technology). EEG recordings were performed when the animals were 3 months old.

### Data Analysis and Statistics

The data analysis was performed using *Luna*, a software package for processing sleep EEG signals, and Python. The recorded EEG signals were segmented into 10-second episodes, which were further categorized into different stages, including wake, NREM, and REM, using the Light Gradient Boosting Machine (LightGBM) algorithm (**Fig. 4A**) as we previously published (*39, 101*).

The hypnogram were used to assess macro sleep structures (**Fig. 3E, F, G**). First, the time spent in each stage was calculated. For bout analysis, the lengths of NREM, REM, and wake bouts were calculated, and the mean bout length and its variation were used to assess fragmentation. Fragmentation was further analyzed by dividing the hypnogram into sleep and non-sleep periods (windows), where brief awakenings lasting less than 5 minutes were still considered part of the sleep window. The number of sleep windows during the light and dark cycles and the frequency of awakenings within these windows were quantified. Additional metrics, such as latency to sleep and NREM length as function of time, were also calculated.

The Welch method was employed to perform the spectral analysis. The power values were normalized with respect to the total power of interest for statistical analysis and subsequently converted to the decibel scale for visualization. To investigate rhythmic changes in power, we computed hourly power values in each frequency band and fitted the cosinor function to the data. To assess the extent to which circadian changes explain the variation in power, we calculated R² values based on the group-averaged power trajectories. We also extracted amplitude and phase parameters from fitted curves for each individual recording to further evaluate the strength and phase of 24-hour rhythmic patterns. As a complementary analysis, we assessed whether the phase angles were clustered across animals within each group using the Rayleigh test to measure phase consistency. For connectivity, coherence analysis was performed, which considered both amplitude and phase information of the signal. This method assesses the degree of linear association between two signals, providing insights into their functional connectivity and synchronization. The obtained power spectrum and coherence were then divided into frequency bands for further analysis: slow oscillations (0.5-1 Hz), delta (1-4 Hz), theta (4-8 Hz), alpha (8-12 Hz), sigma (12-15 Hz), beta (15-30 Hz), gamma (30-50 Hz). To compare the periodic component of the power spectrum within the spindle frequency range (7-20 Hz), we used the Fitting Oscillations & One Over F (FOOOF) algorithm (*102*). This method decomposes the power spectrum into periodic (oscillatory) and aperiodic components, allowing for a more direct comparison of oscillatory activity independent of the non-oscillatory background activity. In the analysis of spindle, slow oscillation, and coupling, we employed a wavelet-based analysis toolbox *Luna* as described previously (*40*). Spindles were detected based on the determined threshold of WT animals. We characterized sleep spindle properties including density (# of spindles per minute), amplitude, integrated spindle activity (ISA) (the sum of normalized wavelet coefficients in the spindle interval), and duration. We then determined the relative position of spindles with respect to the SOs for coupling analysis, as illustrated in **Fig. 3D**.

Outliers were excluded before statistical tests using the interquartile range (IQR) method, excluding values that did not fall between Q1 - 2IQR to Q3 + 2IQR. For analyses without repeated measures, one-way ANOVA was used for comparisons among three or more groups. When repeated measurements were present, we employed a linear mixed model, allowing for a more robust evaluation of the effect of the independent variables (*103*). Tukey’s post hoc test was applied for multiple comparisons. Circular statistics were conducted using the Watson-Williams test and Rayleigh test for uniformity. Statistical significance between genotypes and frequency bands was determined using a threshold of p < 0.05.

### Bulk RNA Sequencing and Data Analysis

Total RNA was extracted from tissues preserved in RNAlater using the Qiagen RNeasy Plus Universal Mini Kit, following the manufacturer’s protocol (Qiagen, Hilden, Germany). RNA quantity was assessed using a Qubit 2.0 Fluorometer (Thermo Fisher Scientific, Waltham, MA, USA), and integrity was evaluated using a TapeStation (Agilent Technologies, Palo Alto, CA, USA). Only samples with an RNA Integrity Number (RIN) greater than 8 were used for library preparation.

RNA sequencing libraries were prepared using the NEBNext Ultra II RNA Library Prep Kit for Illumina, according to the manufacturer’s instructions (New England Biolabs, Ipswich, MA, USA). Briefly, mRNA was enriched using oligo(dT) beads and fragmented at 94°C for 15 minutes. First and second-strand cDNA synthesis followed, after which cDNA fragments were end-repaired and adenylated at the 3’ ends. Universal adapters were ligated to the cDNA, followed by indexing and PCR enrichment with limited cycles. The sequencing libraries were profiled using the Agilent TapeStation (Agilent Technologies) and quantified using a Qubit 2.0 Fluorometer and quantitative PCR (KAPA Biosystems, Wilmington, MA, USA). Libraries were sequenced on the Illumina HiSeq 4000 with a 2×150 base paired-end configuration, at least 30M paired-end reads per sample. Image analysis and base calling were performed using the instrument’s control software, and raw sequence data (BCL files) were converted to FASTQ files and de-multiplexed using Illumina’s bcl2fastq 2.20 software, allowing for one mismatch in index sequence identification.

Paired-end FASTQ files were aligned to the mouse genome (GRCm39, GENCODE M34) and quantified using Salmon v0.14.1 (*104*). Transcript-level data were aggregated to the gene level in R using tximport v1.30.0 (*105*). To identify and account for unmodeled sources of variation in the RNA-seq data, such as batch effects, we implemented Surrogate Variable Analysis (SVA) (*106*). Using the sva R package, we estimated the number of significant surrogate variables (SVs) from the normalized count data. Two or three significant surrogate variables were identified and subsequently incorporated as covariates into the main DESeq2 (v1.34.0) model (*107*). The differential expression analysis was then performed using the final design formula: ∼ SV1 + SV2 + genotype + period + genotype:period. This approach allows the model to account for hidden variation during statistical testing. For data visualization, such as in Principal Component Analysis (PCA) plots, a batch-corrected expression matrix was generated. The count data was first transformed using a variance-stabilizing transformation (VST) from the DESeq2 package. Subsequently, the variation attributable to the two surrogate variables was removed from the transformed data using the removeBatchEffect function from the limma R package (*108*). It is important to note that this batch-corrected data matrix was used exclusively for visualization, while all statistical testing for differential expression was performed on the original counts using the DESeq2 model described above. Differential gene expression analysis was performed with DESeq2, identifying differentially expressed genes (DEGs) with an adjusted p-value < 0.05. Gene set enrichment analysis (GSEA) was conducted using fgsea v1.28.0 (*59*), utilizing gene ontology (GO) terms from the Mouse Molecular Signatures Database (MSigDB) v2023.2 (*109*). Additional gene sets related to activity-regulated genes, rhythmic genes, glucocorticoid genes, and sleep deprivation-regulated genes were curated from literature or obtained in house (Table S3). Transcription factor activity was inferred using decoupleR v2.12.0 (*65*).

### Nuclei Extraction, sn RNA-seq Library Preparation and Analysis

A single hemisphere of each brain was dissociated in 2 ml of prechilled EZ lysis buffer (Nuc-101, Millipore Sigma) via douncing using tissue grinder sets (D8938, Millipore Sigma) and allowed to incubate for 5 minutes on ice. The lysate was passed through a prewet 30 micron filter (04-0042-2316, Sysmex), and 1.5 ml of MST wash Buffer (10 mM Tris, 146 mM NaCl, 1 mM CaCl2, 21 mM MgCl2, 1% BSA, 0.02% Tween-20, 40 U/ml RNase Inhibitor (2313, Takara)), was used to collect the remaining nuclei from the mortar. The samples were centrifuged for 5 minutes at 500 g and 4°C before removal of the supernatant and completing a second wash. The nuclei were resuspended and counted with AOPI (CS2-0106, Revvity) on a Cellometer K2 (CMT-K2-MX-150, Revvity). Samples were loaded on the 10x Chromium Next GEM Single Cell 3’ Kit V3.1 (Dual Index, PN-1000268, 10x Genomics). Library construction followed the 10x guidelines (CG000315 RevF), and sequencing was performed with a NovaSeq S2 flowcell (Illumina) with 28 bases for read 1, 90 bases for read 2, 10 bases for index read 1 and 10 bases for index read 2, aiming for the recommended read depth for each sample.

FASTQ files were processed with Cell Ranger v7.1.0 (*110*), setting the --expect-cells argument based on the expected number of nuclei loaded into the chip, --include-introns=true, --chemistry=SC3Pv3, and using the 10x refdata-gex-mm10-2020-A reference. For each sample, DropletUtils v.1.14.2 (*111*) was used to load the associated molecule_info.h5 into R v4.1.3 (*112*) with the downsampleReads command, downsampling all samples to the same number of reads per cell based on the expected number of cells. A count matrix was produced with emptyDropsCellRanger, with n.expected.cells set to the expected number of cells, umi.min=300, and cand.max.n=20000, returning an FDR score per droplet. All droplets with FDR > 0.1 were excluded, and the rownames of the count matrix were changed to gene names. A Seurat v4.0.0 (*113*) object was created in R v4.0.3 from this count matrix with CreateSeuratObject and normalized with NormalizeData. Cells with < 250 genes expressed were removed. Variable genes were found with FindVariableFeatures, scaled with ScaleData, and PCA was performed with RunPCA with npcs=60. Dimensionality reduction with scGBM v0.1.0 (*114*) was performed on the count data for the variable genes, with M=20 and subset=50000. This dimensionality reduction was added to the Seurat object, and clustering was performed on it with FindNeighbors and FindClusters, while the UMAP was performed on it using RunUMAP. Doublet scores were calculated on each sample with scds v1.6.0 (*115*) Azimuth v0.3.2 (*116*) was used to get an initial cell type labelling with single-nucleus RNA-seq data from the Allen brain atlas whole cortex and hippocampus 10x Chromium dataset (*117*) as the reference using the subclass_label. Cell level QC was calculated for each sample with our single cell QC tool (https://github.com/seanken/CellLevel_QC) and added to the metadata for the Seurat object, as was the percent of UMIs from the mitochondria. Clusters with an average scds score > 0.7 or an average % intronic reads < 50% were removed. scGBM, clustering, and the UMAP were recalculated on this filtered object using the same settings as previously. Clusters were annotated by cell type using the annotations from Azimuth, scores from scds, and manual curation by marker genes. We removed contaminating cells from other brain regions (Meis2+ inhibitory clusters, hippocampal neurons, clusters labelled as Car3, and unannotated neurons) and doublets. Finally, cells with > 2% mitochondrial reads, > 5% ribosomal proteins, > 25,000 UMIs, or < 25% intronic reads were removed, to generate the final Seurat object.

Changes in cell type composition were calculated with the propeller.ttest command (with robust=TRUE and trend=FALSE) in the scater package v.1.18.5 (*118*) using the asin transformation. For differential expression between conditions, a pseudobulk-based approach was used. More specifically, for each cell type the per sample pseudobulk (based on the sum of all cells in that sample) was calculated using a slightly modified version of the code the AverageExpression command in Seurat. Samples with < 20 cells were excluded, and cell types with <2 samples per condition after filtering were not tested. Using edgeR v.3.32.1 (*119*) a DGEList object was created, and genes were filtered using the filterByExpr command with default parameters. Size factors were calculated with calcNormFactors and dispersion parameters were estimated with estimateDisp, using default parameters. The model was fit with glmFit, and the LRT test was performed with glmLRT for each coefficient of interest. P-values were rescaled with fdrtool v.1.2.16 (*120*) (using the z score calculated as -sign(res$logFC)*qnorm(res$PValue/2)), and FDR-corrected p-values were calculated with p.adjust. Enrichment analysis was performed with fgsea v.1.16 (*59*) using the signed log p-value as the score. GO terms for the enrichment analysis were extracted from AnnotationDbi v1.52 (*121*) Finally, TRADE v.0.1.0 (*55*) was used to measure the size of the transcriptomic change. More specifically, first AggregateExpression with slot=“counts” was used to calculate a per sample pseudobulk for each cell type, and DESeq2 v1.30.1 (*107*) was used to calculate DE genes between conditions (using DESeqDataSetFromMatrix to create the DESeq object, the DESeq command to fit the model, and the results command to get the DE output). The TRADE score was then calculated with the TRADE command, with mode = “univariate”. The per mouse condition labels were permuted and the differential expression and TRADE score were calculated on this permuted data. This was repeated 100 times and a permutation p-value was calculated for each cell type.

### Biochemical Fractionation and Quantitative Proteomics

Mouse prefrontal cortex and hippocampus tissue was collected and homogenized in a 20X ice-cold homogenization solution composed of 320 mM sucrose, 10 mM HEPES (pH 7.4), 1 mM EDTA, 5 mM sodium pyrophosphate, 1 mM sodium vanadate (Na□VO□), and a protease inhibitor cocktail (Roche, Switzerland). Following homogenization, the samples were centrifuged at 800 × g for 10 minutes to separate cellular debris. The resulting supernatant (designated as S1) was subjected to a second centrifugation at 16,000 × g for 20 minutes at 4°C to isolate the P2 fraction. The P2 fraction was then solubilized in a buffer containing 1% Triton X-100, 50 mM Tris (pH 7.4), 150 mM NaCl, 1 mM EDTA, and supplemented with protease and phosphatase inhibitors. This mixture was incubated on ice for 20 minutes to ensure complete solubilization. After incubation, the samples were centrifuged at maximum speed (benchtop) for 30 minutes at 4°C. The supernatant obtained from this centrifugation was designated as the synaptic fraction.

For quantitative proteomic analysis, the synaptic fractions from each sample were reduced, alkylated, and digested with trypsin (1:100 enzyme to protein ratio) overnight at 37°C. Tryptic peptides were labeled using a TMTpro 16plex labeling kit (Thermo Fisher Scientific), according to the manufacturer’s instructions. Labeled peptides were pooled and desalted using a C18 spin column (Thermo Pierce), followed by vacuum drying. TMT-labeled peptides were separated by reverse-phase high-performance liquid chromatography (HPLC) using a Thermo Ultimate 3000 system equipped with a Thermo PepMap RSLC C18 column (2 μm, 75 μm × 50 cm). The mobile phases were solvent A (0.1% formic acid in water) and solvent B (0.1% formic acid in acetonitrile). Peptides were eluted over a 140-minute gradient with the following conditions: 1% B for 0–10 minutes at 300 nL/min, 1% B for 10–15 minutes with a flow rate decreasing from 300 nL/min to 200 nL/min, 1–5% B for 15–20 minutes at 200 nL/min, 5–25% B from 20–104.8 minutes at 200 nL/min, 25–35% B from 104.8–112 minutes at 200 nL/min, 35–80% B from 112–115.5 minutes at 200 nL/min, 80% B for 115.5–120 minutes at 200 nL/min, and 80–1% B from 120–140 minutes at 200 nL/min. Peptides were analyzed using an Orbitrap Exploris 480 mass spectrometer (Thermo Fisher Scientific) in data-dependent acquisition (DDA) mode. Full MS scans were acquired at a resolution of 60,000 across a mass range of 450–1600 m/z with a maximum injection time of 50 ms. MS/MS spectra were collected at a resolution of 45,000 with a normalized collision energy (NCE) of 32 and a dynamic exclusion time of 30 seconds. Raw data files were processed using Proteome Discoverer software (version 2.4, Thermo Fisher Scientific). Tandem mass spectra were searched against a mouse reference database (GRCm39) and an in-house contaminant database using the Sequest HT algorithm. Search parameters included a precursor mass tolerance of 10 ppm, fragment mass tolerance of 0.05 Da, and up to 2 missed cleavages allowed for trypsin. Fixed modifications included carbamidomethylation of cysteine and TMTpro modification on lysine residues and peptide N-termini. Variable modifications included methionine oxidation, N-terminal acetylation, and methionine loss.

### Home cage monitoring

*Cacna1g*+/- (HT) male mice and their wild-type littermates (WT) were individually housed in DVC Digital Ventilated Cages (Tecniplast, Buguggiate, Italy) equipped with electromagnetic sensors embedded in the rack infrastructure. Continuous locomotor activity monitoring commenced on postnatal day 28 and extended for 60 days. The electromagnetic sensors captured real-time movement patterns by detecting changes in the electromagnetic field around the mice, enabling precise and non-invasive tracking of locomotion. Activity data were aggregated into 5-minute interval bins and exported for downstream analysis.

### Motion sequencing (MoSeq)

Cacna1g WT, HT, and KO mice were individually tested in a circular open-field arena (diameter: 18″, wall height: 15″) under dim red light conditions. All recordings were conducted during the dark period between 8:00 PM and 11:00 PM to align with the animals’ active phase. Mice were habituated to the experimental room for 10 minutes in their home cage before being placed in the center of the enclosure. Spontaneous behavior was recorded for 30 minutes using a high-resolution overhead camera. No food, water, bedding, or enrichment was present in the arena during the recording period. The enclosure was cleaned with 70% ethanol between sessions to minimize olfactory cues.

Behavioral videos were processed using the Motion Sequencing (MoSeq) pipeline v1.3.0 (*122*) to extract three-dimensional representations of body posture and movement. Background subtraction and body-centered alignment were applied to raw video frames to track the mouse’s body position. A Gaussian smoothed representation of body posture was then used as input for an autoregressive hidden Markov model (AR-HMM), which identified a set of discrete, recurring syllables corresponding to stereotyped behavioral motifs. Syllable usage was quantified as the relative frequency of each syllable across genotypes. Log□ fold changes in syllable frequency were calculated for HT vs. WT and KO vs. WT comparisons. Linear discriminant analysis (LDA) was used for dimension reduction to check for the presence of clusters among groups based on movement features, including centroid position, velocity, and posture. Heatmaps were generated to visualize spatial occupancy during the recording period. Syllable classification was achieved by an experimenter visualizing individual syllables in their movie form and assigning a label empirically. Statistical comparisons of syllable usage were performed using Kruskal-Wallis tests, followed by post hoc pairwise Wilcoxon rank-sum tests where appropriate. Data are presented as mean ± 95% confidence interval (CI), with p < 0.05 considered statistically significant.

